# Biogenesis of mature piRNAs in *Caenorhabditis elegans* is dependent on orchestration between multiple ribonucleases and PRG-1

**DOI:** 10.1101/2021.12.29.474424

**Authors:** Aniruddha Samajdar, Tamoghna Chowdhury, Saibal Chatterjee

## Abstract

Piwi-interacting RNAs (piRNAs) are an animal-specific class of germline-enriched small non-coding RNAs that shape transcriptome, as well as ensure genomic integrity and fertility by regulating transposons and other selfish genetic elements. In *Caenorhabditis elegans* mature piRNAs are 21-nucleotides long, begin with a monophosphorylated uridine, and they associate with PRG-1 to form piRISCs that scan the transcriptome for ‘non-self’ sequences. However, these piRNAs are born as longer 5’-capped transcripts, where PARN-1, a 3’-5’ exoribonuclease, contributes to the formation of the mature 3’-end. But, till date, the 5’-processing events remain elusive. We demonstrate that the recently identified endoribonuclease activity of XRN-2 is involved in the processing of the 5’-end of precursor piRNAs in worms. Depletion of XRN-2 results in reduced mature piRNA levels, with concomitant increase in levels of the 5’-capped precursors. We also reveal that the piRNAs born as longer precursor molecules (≥60 nt), prior to 5’-end processing, undergo ENDU-1-mediated endoribonucleolytic processing of their 3’-ends. Our *in vitro* RNA-protein interaction studies unravel the mechanistic interactions between XRN-2 and PRG-1 towards the formation of mature 5’-ends of piRNAs. *In vivo* experiments employing *prg-1* mutant worms indicate that XRN-2 has the potential to perform clearance of precursors that are not bound and protected by PRG-1. Finally, we also demonstrate that XRN-2 is not only important for the generation of mature piRNAs and piRNA-dependent endo-siRNAs, but through yet unknown pathways, it also affects piRNA-independent endo-siRNAs that shape transcriptome, as well as contribute to genomic integrity via regulation of transposable elements.

## Introduction

Piwi-interacting RNAs (piRNAs) are an animal-specific class of germline-enriched small non-coding RNAs that protect animal germlines from transposons and other selfish genetic elements, and thereby, ensure genomic integrity and fertility (**1, 2**). They remain associated with the PIWI clade of Argonaute proteins and repress their targets both co-transcriptionally and post-transcriptionally (**2**). However, the exact molecular mechanisms by which piRNAs carry out their functions do indeed vary amongst different species.

In *Caenorhabditis elegans* (*C. elegans*), mature piRNAs are predominantly 21-nucleotides (nt) long with a strong bias for a 5’-monophosphorylated uracil as the first nucleotide (thus they are also known as 21U-RNAs), and a 2’-O-methylated 3’-residue (**3-5**). They associate with PRG-1, one of the two PIWI clade Argonaute proteins encoded by the *C. elegans* genome, and form the piRNA-induced silencing complex (piRISC). 21U-RNA-PRG-1 complexes scan for foreign sequences, while allowing significantly high level of mismatched pairing with the target transcripts (**6, 7**). Such relaxed sequence complementarity between the 21U-RNA and the target RNA poses the potential risk that piRNAs might target germline-expressed self-transcripts and silence them. Intriguingly, in order to avoid wrongful silencing of *bona fide* germline transcripts, *C. elegans* employs CSR-1-22G-RNA pathway to protect germline-expressed self-transcripts from 21U-RNA pathway-mediated silencing (**7, 8**). The present knowledge indicates that silencing or licensing of a target-transcript will depend on the outcome of a competition between PRG-1 and CSR-1 targeting. If PRG-1 targeting dominates, a transcript is flagged as non-self and targeted for silencing. Conversely, if CSR-1 targeting prevails, the transcript is recognized as self and allowed to express. Upon target recognition, piRISCs do not directly cleave their target transcripts. Instead, they recruit RNA-dependent RNA polymerases (RdRPs) such as RRF-1 and EGO-1 at the target sites that initiate the *de novo* synthesis of 22-nt long endogenous small interfering RNAs (endo-siRNAs) by using the target transcripts as template. These endo-siRNAs show a strong bias for 5’ guanosine residue (also referred as “22G-RNAs”, **references 7-13**). 22G-RNAs are then loaded onto worm-specific WAGO clade of Argonaute proteins (WAGO-1, WAGO-9/ HRDE-1, and WAGO-10) to direct the target silencing, both post-transcriptionally and co-transcriptionally (**6-9**). 22G-RNAs can be transmitted transgenerationally and as a result, silencing of the targets by piRNA pathway can become stable and maintained across several generations, even in the absence of active 21U-RNA-PRG-1 (**7**). Accordingly, *prg-1* mutant worms that lack mature piRNAs, do not become sterile immediately, rather, they show gradual decline in the brood size across generations and eventually become completely sterile after several generations (**9-12**).

Given the functional importance of 21U-RNAs in driving a very powerful silencing response in the germ cells of *C. elegans*, the biogenesis of 21U-RNAs is of paramount importance for the continuity of its life cycle. But, little is known about the chronological events leading to the formation of a mature piRNA. A vast majority of the worm piRNA genes (∼15,000) are located on two discrete clusters of 2.5 Mb (4.5-7 Mb) and 3.7 Mb (13.5-17.2 Mb) on chromosome IV, and primarily reside in introns and intergenic regions (**3, 5**). Each of these piRNA genes located in these two clusters is characterized by the presence of a conserved ‘CTGTTTCA’ motif (Ruby motif) at its promoter region, and collectively these are known as motif-dependent piRNAs or type I piRNAs (**3, 5, 14, 15**). Another subset of low-abundance piRNAs (∼5% of the total piRNA population) do emerge from transcription start sites (TSSs) of protein-coding genes flanked by YRNT motifs (motif-independent or type II piRNAs, **references 14-15**).

Both types of piRNAs are largely transcribed as short 27-40 nt capped RNA precursors by RNA polymerase II (**14-16**). The precursor-transcription initiates in a way so that pre-piRNAs possess two extra nucleotides upstream of the 5’-end of the corresponding mature 21U-RNAs, i.e.; 5’-ends of the precursors are extended beyond the mature sequence by 3-nt including the cap. However, the length of their 3’-extensions beyond the 21^st^ nucleotide of the mature sequences vary among different 21U-RNA species. Therefore, to generate a mature 21U-RNA, the 5’ cap and first two nucleotides from the 5’-end must be removed, and the precursor sequences at the 3’-end would also need to be clipped off. It has previously been shown that PARN-1, a conserved 3’-5’ exoribonuclease, contributes to the 3’-end processing of the precursors (**17**). *parn-1* mutant worms accumulate populations of 5’-end processed piRNAs, where their 3’-ends are extended by ∼2-4 nt beyond that of the mature sequence. However, there are some piRNAs with relatively longer precursor size (40-100 nt; **references 14, 16, 18**). In such cases, PARN-1 mediated processing event takes place downstream of a nucleolytic event by an unknown enzyme, which removes a longer stretch of nucleotides at the 3’-end of the precursor, and thus leaving 2-4 extra nucleotides beyond the mature sequence that needs to be acted upon by PARN-1. This indicates that PARN-1 is not the solo player in the 3’-end processing event of *C. elegans* piRNAs. Moreover, the exact molecular mechanism underlying the 5’-end processing remains unknown. Hypothetically, maturation of the 5’-end can be achieved by the function of a decapping enzyme, followed by a 5’-3’ exoribonucleolytic activity. Alternatively, an endoribonuclease can alone cleave the excess nucleotides along with the 5’-cap to generate the mature 5’-end. PETISCO/ PICS, a recently identified multi-protein complex, has been implicated in 21U-RNA biogenesis and holds potential to bind the 5’-cap and 5’-phosphate of RNA (**19-21**). But, neither this complex contains any decapping enzyme nor does it host any ribonuclease(s). This clearly suggests that there must be additional players, who carry out critical functions towards the processing of the 5’-end.

Here, employing a combination of genetic and biochemical approaches, we demonstrate that the recently identified endoribonuclease activity of XRN-2 is involved in the processing of the 5’-end of precursor piRNAs in *C. elegans*. Depletion of XRN-2 leads to a reduction in mature piRNA levels, with concomitant increase in the levels of 5’-unprocessed ∼30-nt long precursors. We report that in case of 21UR-1, which is transcribed as a longer precursor molecule (∼60-nt), the 5’-end processing happens downstream of an endoribonucleolytic processing of the 3’-end of the transcript, which is mediated by ENDU-1. Here, we also present a partial reconstitution of the precursor piRNA processing *in vitro*, which allows us to appreciate the mechanistic interactions between XRN-2 and PRG-1 towards the formation of mature 5’-ends of piRNAs. Furthermore, *in vivo* experiments employing *prg-1* mutant worms indicate that XRN-2 has the potential to perform clearance of precursors that are not bound and protected by PRG-1. Finally, we also demonstrate that XRN-2 is not only important for the generation of mature piRNAs and piRNA-dependent endo-siRNAs, but it also affects piRNA-independent endo-siRNAs that shape transcriptome, as well as contribute to genomic integrity by modulating the homeostasis of transposable elements.

## Results

### XRN-2 is important for piRNA-mediated repression of target gene

In order to identify the factor(s) involved in the 5’-end processing of precursor 21U-RNAs, we performed an RNAi-based genetic screen employing a *C. elegans* piRNA reporter strain **(9)**. This reporter strain hosts a single copy of a transgene construct on its chromosome II. The transgene comprises the promoter sequence of *mex-5*, followed by the coding sequence of *gfp* fused to the coding sequence of a germline expressed histone (*his-58)* and the 3’ UTR of *tbb-2*. Upstream of the 3’ UTR, it harbors the target site of an endogenous piRNA (21UR-1). Thus, in the wild-type scenario, GFP gets efficiently silenced in the germline of the reporter strain by piRISC, whereas disruption of gene(s) involved in the piRNA pathway (e.g. *prg-1*) results in the derepression of GFP.

We examined all the known *C. elegans* decapping enzymes and 5’-3’ exoribonucleases for their ability to de-silence GFP upon depletion through RNAi by feeding, initiated on synchronized L1 stage larvae at 20°C. We specifically chose this temperature because it has been reported that the wild-type animals show reduction in the mature 21U-RNA levels at an elevated temperature (**≥** 24°C, **3, 20**). Amongst the candidate genes, depletion of XRN-2 and DOM-3 led to the GFP desilencing in the germline of 66% (± 4.1, n = 50, N = 3) and 18.46% (± 0.6, n = 50, N = 3) of the experimental worms, respectively, whereas depletion of PRG-1 (positive control) led to desilencing in 74% (± 2.5, n = 50, N = 3) of the experimental worms (**Figure 1A, Supplementary Table 1**). This outcome indicated to a possible role of XRN-2 and DOM-3 in piRNA metabolism in *C. elegans*. Accordingly, the levels of the primary transcript as well as the spliced mRNA corresponding to the reporter construct were upregulated upon XRN-2 and DOM-3 depletion, consistent with co-transcriptional or a combination of post- and co-transcriptional regulation of targets by the piRNA pathway (**Figure 1B**).

**Figure 1.**
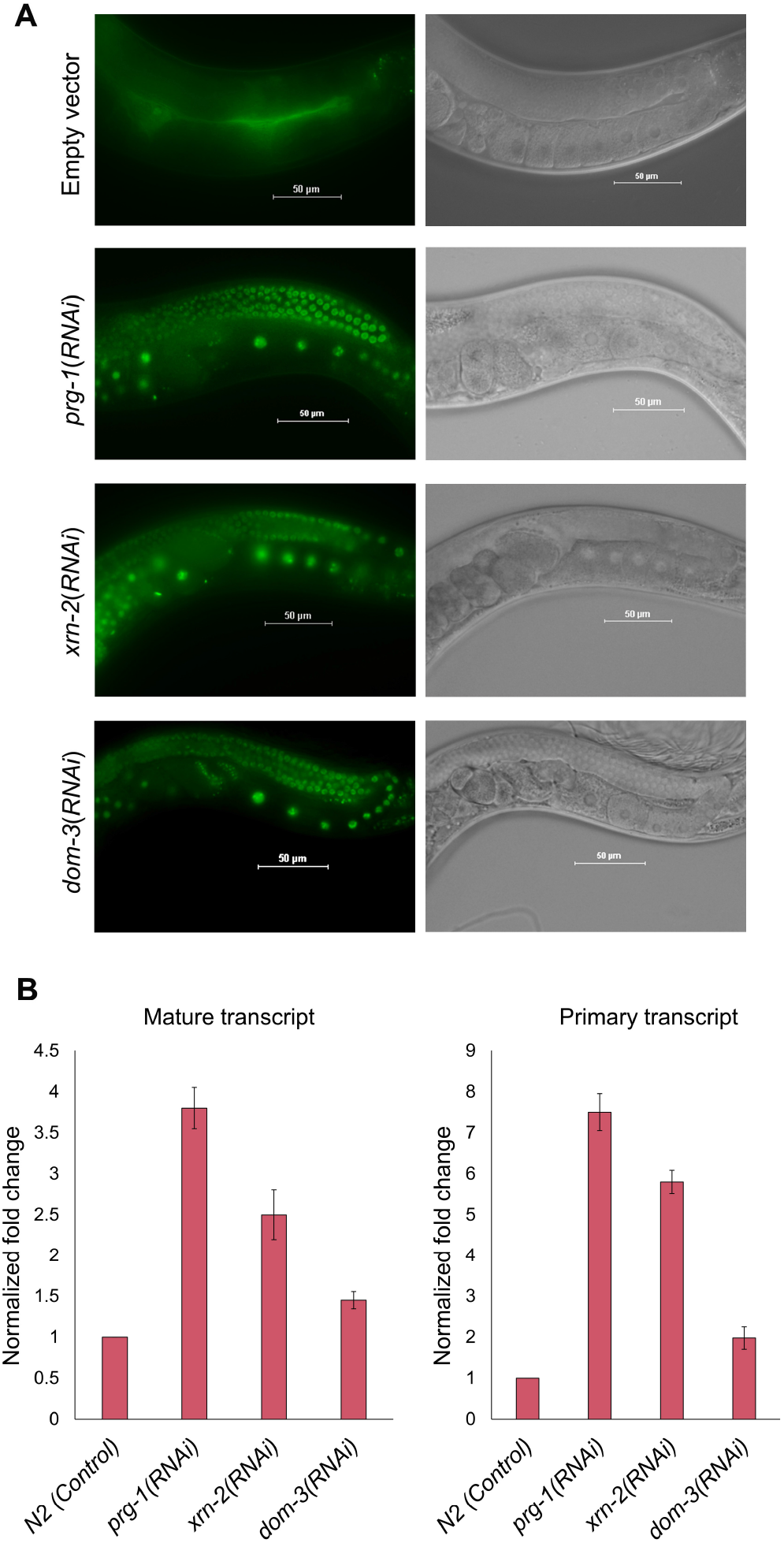
*xrn-2*(*RNAi*) results in the derepression of a piRNA (21UR-1) reporter transgene. **A.** Germline expression of GFP-reporter gene (GFP-H2B) remains repressed by 21UR-1 in wild-type control worms (empty vector), but gets derepressed upon depletion of PRG-1 (positive control) and XRN-2, DOM-3 (candidate genes). Epifluorescence images of the worm germline are depicted on the left, and corresponding DIC images on the right. **B.** RT-qPCR quantification (n = 3; means ± SEM) of the primary transcript and the spliced mature mRNA corresponding to the piRNA reporter construct from the four samples depicted in panel A.

### Depletion of XRN-2 results in the diminishment of mature piRNAs and deregulation of specific targets

Based on the current knowledge about XRN-2’s varied roles at the level of transcriptional and post-transcriptional events (**22-25**), its depletion might impact 21U-RNA-PRG-1 pathway at a single or multiple levels. Given the roles of XRN-2 in transcription termination and other transcriptional events (**22, 23**), it’s depletion might deregulate the synthesis of precursor 21U-RNAs, which in turn, might affect mature 21U-RNA levels. On the other hand, owing to its dual ribonucleolytic activities (**24**), XRN-2 could be required for the processing of precursor 21U-RNAs into mature 21U-RNAs, and thus, its loss might result in the accumulation of some precursor forms. Additionally, it is also possible that XRN-2 might not be involved with the transcription or processing of 21U-RNAs at all, rather it might directly or indirectly (owing to its role in rRNA biogenesis and translation, **reference 22**) affect subunits of piRISC that may modulate the efficacy for target binding and activity of the effector complex. Notably, in our experimental samples the level of PRG-1 remained unchanged, while XRN-2 was depleted (**Supplementary Figure 1A**). XRN-2 might also influence the RdRP system and/or 22G-RNA production, through which piRISCs extend their function. Under the two aforementioned circumstances, depletion of XRN-2 might not result in any change in the precursor as well as mature 21U-RNA levels, but would still exert an effect on target silencing. Of note, to exclude the possible indirect effects of deregulated rRNA metabolism, we standardized the duration and extent of *xrn-2(RNAi)* in a way that at the chosen time point for performing molecular analyses (young adult), there was very little or no effect on rRNA levels (**Supplementary Figure 1B**), yet the reporter strain showed derepression of GFP.

Therefore, in order to understand the molecular role of XRN-2 in piRNA metabolism, we decided to check the levels of precursor and mature 21U-RNAs, upon its depletion. We performed northern blotting, where we probed for 21UR-1, one of the most abundant piRNAs. Notably, most piRNAs have very low GC-content, and only a few piRNAs have optimal sequence for detection through northern probing. Total RNA extracted from XRN-2 depleted young-adult worms revealed three distinct species of 21UR-1, a 21-nt long mature form and two precursor forms of ∼30-nt and ∼60-nt length (**Figure 2A, Supplementary Figure 1C**). This ∼60-nt band corresponds to the expected RNA Pol II transcript, and also got diminished upon depletion of the known piRNA transcription factor UNC-130 (henceforth, referred to as ‘60-nt precursor-transcript’) (**Supplementary Figure 1D**), and the ∼30-nt band represents a processing intermediate (henceforth, referred to as ‘30-nt intermediate-precursor’). Interestingly, in *xrn-2*(*RNAi*) samples, levels of mature 21UR-1 got dramatically decreased, with a concomitant increase in the level of the 30-nt intermediate-precursor molecules (**Figure 2A, 2B**). We observed that these 30-nt intermediate-precursors in XRN-2 depleted worms were 5’-capped as they were unaffected by the treatment of a 5’-3’ terminator nuclease, which again confirmed also suggested that XRN-2 is capable of processing a 5’-capped substrate (**Figure 2C**). Moreover, we observed that this 30-nt intermediate-precursor distinctly migrates slower than the precursor band that accumulates upon depletion of PARN-1 (**17**), which suggested that 3’-end processing of 21UR-1 by PARN-1 might be dependent on 5’-end processing by XRN-2 (**Figure 2D**). Notably, the ‘30-nt intermediate-precursors’ that we could detect through RNA sequencing of the experimental samples (subjected to decapping reaction before library generation), possessed two extra nucleotides at their 5’-end, appended to the *bona fide* first uridine residue of the mature piRNA. And for all of them, we also did not detect any template-independent 3’-appended nucleotides (**data not shown**). These observations suggested that the size of the 30-nt intermediate-precursor of 21UR-1 was indeed a derivative of template-dependent transcripts and not template-independent nucleotide additions. Additionally, there was no change in the level of the 60-nt precursor-transcript molecules, compared to that of the control sample (**Figure 2A, 2B**). These results indicated that XRN-2 does not affect the transcription of *21ur-1* gene, rather it is involved in the processing of the 30-nt intermediate-precursor into the corresponding mature form. XRN-2 depletion also exerted a similar kind of effect on the precursor and final product of *21ur-1949*. For 21UR-1949, we could detect a single band corresponding to the mature form in the control scenario, whereas XRN-2 depleted worms failed to generate mature 21UR-1949 and instead accumulated a longer 5’-unprocessed species of ∼30-nt in length (**Figure 2E**). Notably, for 21UR-1949, we were unable to detect any longer precursor similar to that of the 60-nt precursor-transcript of 21UR-1. This could be either due to the inherent instability of such a precursor molecule or due to the absence of such a molecule. Of note, variation in the lengths of the precursor transcripts of different species of 21U-RNAs is well known, where most precursor-transcripts are thought to be ∼30-nt in length (**14, 15, 16, 18**). Together, this result also suggested that irrespective of the length of the precursor-transcripts, XRN-2 is involved in the processing of their 5’-ends. Now, in order to understand the extent to which XRN-2 affects biogenesis of mature 21UR RNAs, we performed small RNA sequencing. We could detect 7493 Type-I 21U RNAs, and ∼60% of those were downregulated (4403) in the XRN-2 depleted sample. These affected 21U RNAs were uniformly distributed across the two loci on chromosome IV (**Figure 2F**). Notably, our sequencing did not capture the ‘30-nt intermediate-precursor’ species as widely as the mature species. Nevertheless, we recorded 146 of these precursors that got upregulated in the experimental sample, for which the corresponding mature forms were downregulated (**Figure 2G, Supplementary Table 2**). We could also detect 133 mature type II piRNAs, among which 78 showed downregulation in the XRN-2 depleted sample. We also detected four type II precursors that showed upregulation in experimental sample, when the corresponding mature forms were downregulated (**data not shown**). Overall, the high-throughput results corroborated with our low-throughput, northern probing-based, observations.

**Figure 2.**
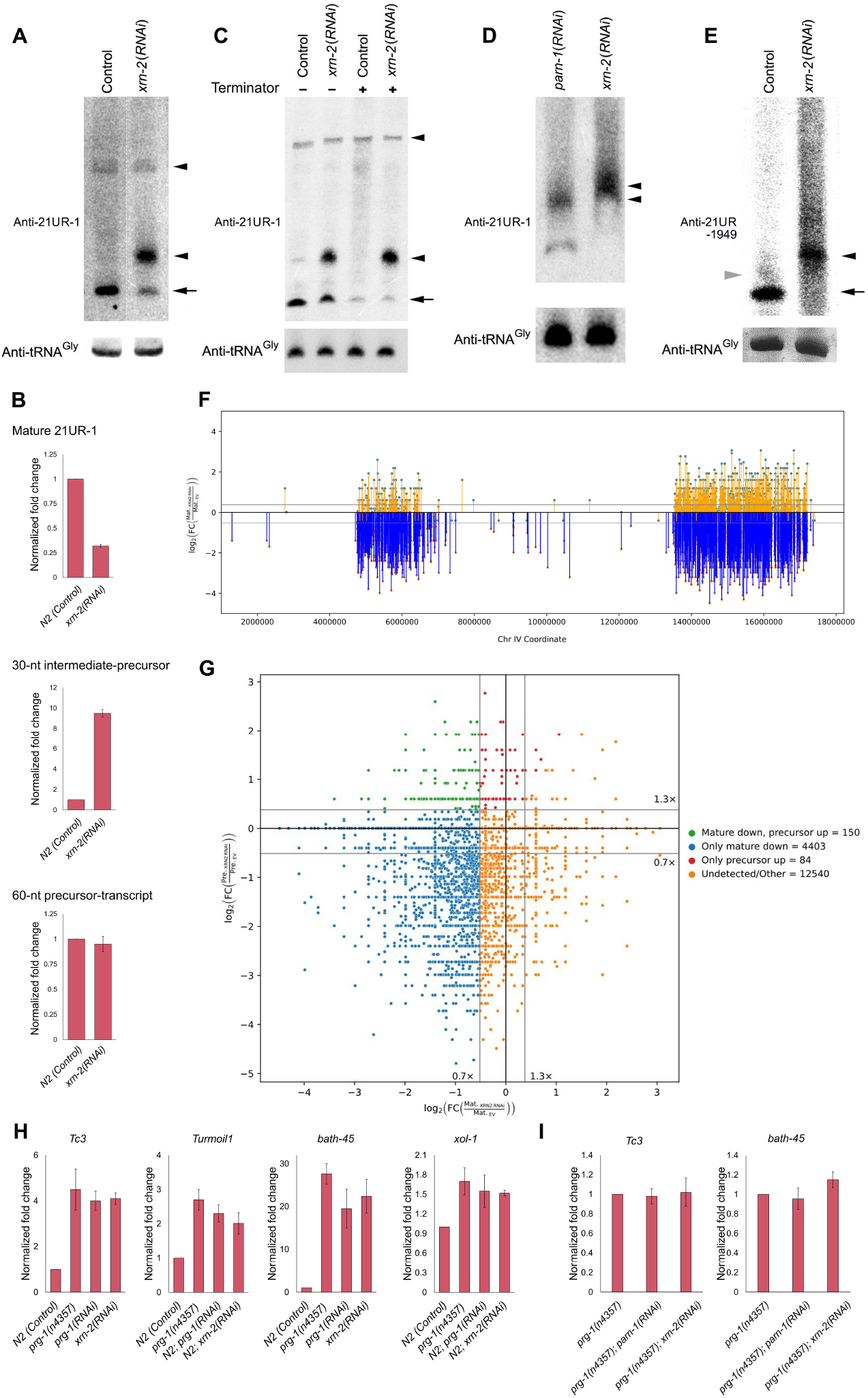
XRN-2 depletion results in diminishment of mature piRNA levels and causes increased expression of targets regulated by the piRNA pathway. **A.** A representative northern image depicting diminished mature 21UR-1 (arrow), hyper-accumulated 30-nt intermediate-precursor (bottom arrowhead), unchanged 60-nt precursor-transcript (top arrowhead) in *xrn-2*(*RNAi*) samples, compared to control (*N2* worms). tRNA^Gly^ served as loading control. Both the lanes are from the same blot. **B.**Graphical representation of the relative levels of the mature and precursor forms of 21UR-1 upon depletion of XRN-2 (n = 3; means ± SEM). **C.**30-nt intermediate-precursors of 21UR-1 (bottom arrowhead) that hyper-accumulate upon XRN-2 depletion are resistant to 5’-monophosphate dependent 5’-3’ exoribonuclease activity of the Terminator enzyme. **D.**The 30-nt intermediate-precursors of 21UR-1 that accumulate upon depletion of XRN-2 show retarded migration compared to the precursors that accumulate upon *parn-1(RNAi)*. The relevant precursor bands are indicated with arrowheads. RNA samples were resolve on a 12% Urea-PAGE. **E.**Accumulation of a longer precursor (black arrowhead) and absence of the mature 21UR-1949 (arrow) in *xrn-2*(*RNAi*) samples. Grey arrowhead on the left indicates to the migration of a 23-nt marker band. **F.**Log_2_fold changes of mature piRNAs of Chromosome IV [*xrn-2*(*RNAi*) vs. Control] on y-axis vs. piRNA transcript start position along Chromosome IV on x-axis. XRN-2 does not show any preference for cluster I or cluster II piRNAs. The cutoffs are 30% upregulated or downregulated, and indicated by grey lines. **G.**Log_2_fold change [*xrn-2*(*RNAi*) vs. Control] of precursor piRNAs from all chromosomes on y-axis vs log_2_fold change [*xrn-2*(*RNAi*) vs. Control] of mature piRNAs from all chromosomes on x-axis. Top left quadrant indicates piRNAs with reduced mature and increased precursor counts upon *xrn-2*(*RNAi*). Grey lines indicate the cutoffs. **H.**RT-qPCR quantification (n = 3; means ± SEM) reveals comparable levels of upregulation of known piRNA targets in samples from *xrn-2*(*RNAi*), *prg-1*(*RNAi*), and *prg-1*mutant worms, compared to control. **I.**RT-qPCR quantification (n = 3; means ± SEM) of known piRNA targets in *prg-1(n4357), prg-1(n4357); parn-1(RNAi)*and *prg-1(n4357); xrn-2(RNAi)*worms. Depletion of XRN-2, as well as the known piRNA pathway component PARN-1, in *prg-1*mutant worms do not change the levels of indicated piRNA targets suggesting their participation in the same genetic pathway.

It has previously been shown that specific transposable elements (e.g. *Tc3, Turmoil1*) and certain endogenous protein-coding genes (e.g. *bath-45* and *xol-1*) get derepressed in *prg-1*mutant worms (**3, 4, 9, 13**). Therefore, derepression of these targets upon XRN-2 depletion would further support XRN-2’s role in 21U-RNA-PRG-1-mediated regulation. Quantification using RT-qPCR revealed that XRN-2 depletion led to the accumulation of mRNAs corresponding to the abovementioned targets to a degree similar to that seen in the *prg-1* mutant worms (**Figure 2H**). To further confirm that the desilencing of the above mentioned targets upon XRN-2 depletion is due to the impairment of piRNA pathway, and not due to effects on some parallel pathways affecting the stability of those targets, we depleted XRN-2 by RNAi in *prg-1*mutant background and checked the levels of those piRNA targets. As a positive control, we also checked their levels in *prg-1; parn-1(RNAi)*worms. If the observed desilencing of *Tc3* and *bath-45* upon XRN-2 knock-down resulted from perturbation of its functions in pathways unrelated to piRNA processing, then *prg-1; xrn-2(RNAi)* animals should show additive/synergistic increase in those transcript levels. However, in contrast to this possibility and consistent with XRN-2’s participation in the same pathway, we found that the transcript levels of both *bath-45* and *Tc3* remained unchanged in *prg-1; xrn-2(RNAi)* animals compared to *prg-1* mutant worms (**Figure 2I**). Furthermore, as expected, their levels also did not change in *prg-1; parn-1(RNAi)* animals compared to *prg-1* mutant worms (**Figure 2I**). These results strongly suggested a role for XRN-2 in piRNA biogenesis. Of note, here we decided not to focus on the role of DOM-3 as its depletion resulted in a modest (∼20%) depletion of mature 21UR-1 and one more piRNA that we could detect through northern probing, and the effects on the known targets that get deregulated in *prg-1* mutant worms were not as much as that observed upon depletion of XRN-2 (**Supplementary Figure 2**). In *Drosophila*, DOM-3’s ortholog (*cutoff*) has been implicated in the transcription of piRNAs from the heterochromatic regions of the genome (**27**), and it is possible that it performs a minor and/or redundant role in piRNA biogenesis in worms.

### Endoribonuclease activity of XRN-2 is responsible for the processing of the 5’-end of the piRNA precursors

Recently, it has been demonstrated that XRN-2 harbors an endoribonuclease activity, and beginning from the 5’-end, it can processively act upon 5’-phosphorylated RNA substrates after every three nucleotides (**24, 25**). Moreover, we noticed that the endoribonuclease activity does not get deterred by a 5’-cap structure (**Supplementary Figure 3**). Such a capability would essentially exclude the explicit requirements of a decapping activity followed by 5’-3’ exoribonucleolysis necessary towards processing of the capped 5’-end of the pre-piRNA. In order to test the possible role of the endoribonuclease activity of XRN-2, we employed a mutant strain with partially perturbed endoribonuclease active site of XRN-2 [*xrn-2*(*PHX25*), **reference 24**]. In expected lines, compared to wild type worms this mutant strain exhibited a diminished level of mature piRNAs (**Figure 3A, 3B**). This reduction in the level of the mature piRNA (21UR-1, 21UR-1949) was functionally relevant as the validated targets (*Tc3, Turmoil1* and *bath-45*) were significantly elevated in the mutant worms, compared to that of the wild type control (**Figure 3C**). But, very unexpectedly, there was very little or no accumulation of the 30-nt intermediate-precursor that accumulated upon depletion of XRN-2 (**Figure 2B, 3A, 3B**). Unchanged level of the 60-nt precursor-transcript excluded the possibility of any adverse effect of this mutant *xrn-2* allele on *21ur-1* transcription, which otherwise could have contributed to a reduced accumulation of this intermediate-precursor. However, it could be possible that the fraction of the 30-nt long intermediate-precursors, which did not get further processed to form the mature 21UR-1 due to perturbation of the endoribonuclease activity of XRN-2 underwent rapid degradation and that precluded their accumulation. It is also possible that such a 30-nt intermediate-precursor, whose 5’-capped end would be bound by the mutant XRN-2 and still remain unprocessed, might possibly be vulnerable from its 3’-end. Accordingly, to investigate whether the piRNA 3’-end processing enzyme PARN-1 plays any role on the 30-nt intermediate-precursor that does not get processed at its 5’-end, we performed depletion of PARN-1 in *xrn-2*(*PHX25*) worms. We observed an accumulation of ∼30-nt long smearish band, similar to the band size of the intermediate-precursor that accumulates upon XRN-2 depletion, which indicated that PARN-1 is indeed responsible for the degradation of the 30-nt long intermediate-precursor that fails to undergo a 5’-end processing (**Figure 3D, 3E**). This suggested a role for PARN-1 as a surveillance factor that prevents the accumulation of a 5’-end unprocessed intermediate-precursor, besides its *bona fide* role in the processing of the 3’-end of the precursor. Additionally, these accumulated smearish bands were 5’-capped as they were unaffected by the treatment of a 5’-3’ terminator nuclease, which again suggested that in the absence of processing of the 5’-capped-end by the endoribonuclease activity of XRN-2, the substrate is made available to the 3’-5’ degradation activity of PARN-1. Notably, the 30-nt intermediate-precursors accumulated upon XRN-2 knockdown do not appear to be the substrates for this PARN-1 mediated surveillance clearance. It indicates to the possibility that endoribonuclease mutant XRN-2 bound to the 5’-end of the 30-nt intermediate-precursor elicits some conformational change probably to PRG-1 that makes the PRG-1 bound-and-protected RNA available for PARN-1 action, which probably doesn’t happen when the 5’-end of the 30-nt intermediate-precursor remains unprocessed due to diminishment of XRN-2. Importantly, upon careful observation, it was clear that the smearish 21UR-1 intermediate-precursor band accumulated upon depletion of PARN-1 in *xrn-2* (*PHX25*) worms indeed comprised of multiple distinct bands, where some of them are larger than the relevant band accumulated upon XRN-2 depletion. Since, these bands are protected from the 5’-end owing to a 5’-cap, they must have been tailed by a 3’-terminal U/A transferase. We performed a screening using candidate approach and identified a gene (*cde-1*) out of ten candidates, whose depletion on top of PARN-1 depletion in *xrn-2(PHX25)* worms resulted in the loss of the tailing event (**data not shown**). Since, many 3’-5’ RNA degradation machineries work downstream of a 3’-tailing event (**28**), our results again suggested that in the absence of appropriate processing of the 3’-end of the intermediate precursor, the precursor may be rendered vulnerable to multiple activities, which in turn might affect generation of functional mature piRNAs. In expected lines, we observed depletion of mature 21UR-1949 accumulation in *xrn-2*(*PHX25*) worms, and without any accumulation of the precursor (**Figure 3D**). Accordingly, the precursor only accumulated when PARN-1 was depleted in the same genetic background, and that too as a smearish product comprising multiple distinct bands (**Figure 3D**). It is interesting to note that the 30-nt intermediate precursors that accumulated upon *xrn-2(RNAi)*, predominantly did not show any template-independent nucleotide addition at their 3’-ends (**data not shown, see materials and methods**), and it indicates to the possibility that XRN-2 might directly or indirectly facilitate the recruitment of components of the relevant 3’-tailing machinery at those unprocessed 3’-ends.

**Figure 3.**
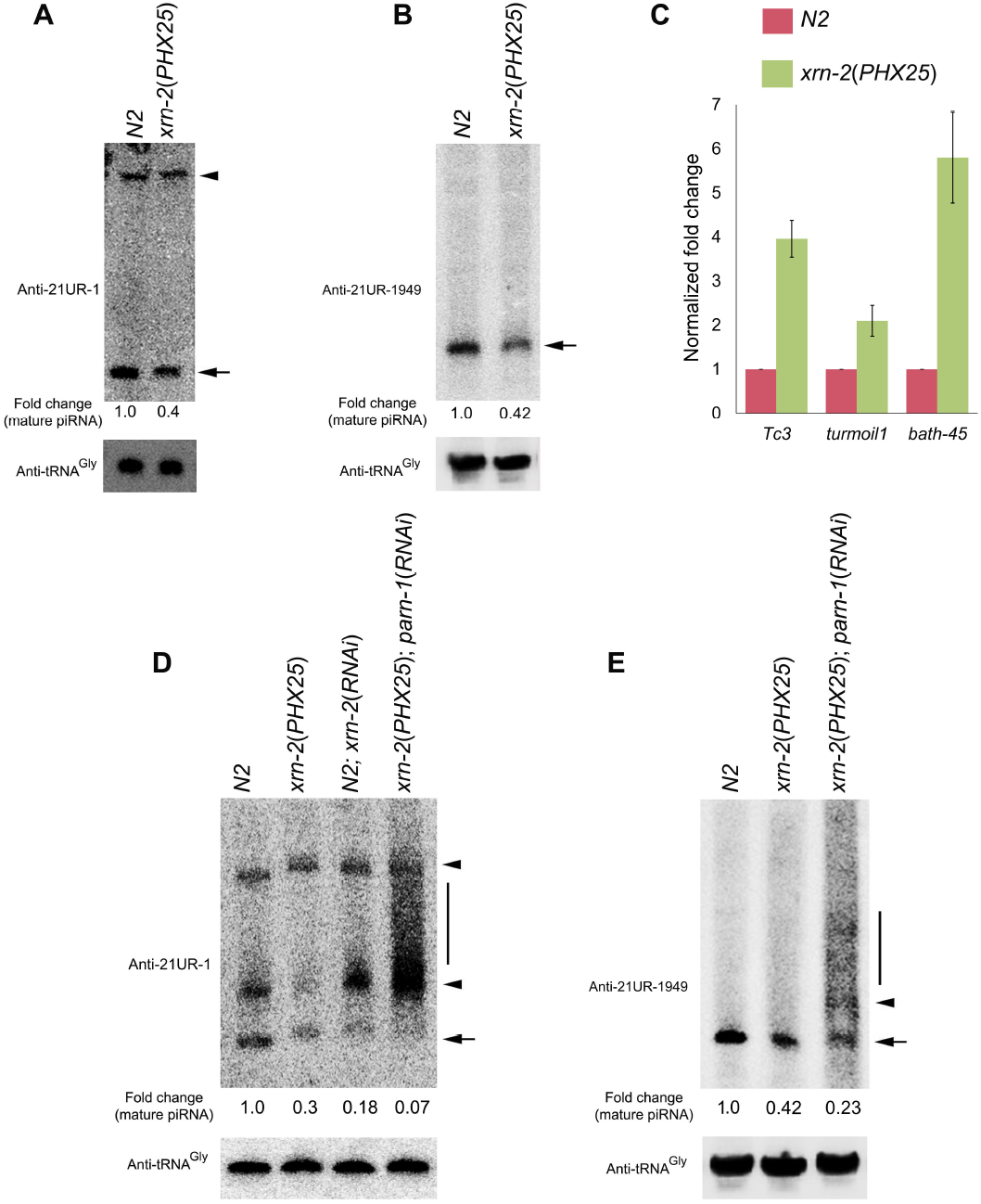
Endoribonuclease activity of XRN-2 is responsible for the processing of the 5’-ends of the piRNA precursors. **A.**A representative northern blot analysis revealing that mature 21UR-1 (arrow) gets depleted in worms mutant for the endoribonuclease activity of XRN-2 [*xrn-2*(*PHX25*)] without any change in the level of the 60-nt precursor-transcript (arrowhead). tRNA^Gly^served as loading control. **B.**Depletion of mature 21UR-1949 (arrow) in *xrn-2*(*PHX25*) worms. **C.**Increased expression of known piRNA targets, as indicated, in *xrn-2*(*PHX25*) worms. **D.**Depletion of PARN-1 results in the accumulation of the 30-nt intermediate-precursor (bottom arrowhead) of 21UR-1 in *xrn-2*(*PHX25*) worms. 5’-capped (data not shown) and smearish nature (indicated by a vertical bar) of the accumulated band suggests 3’-tailing. tRNA^Gly^served as loading control. **E.**21UR-1949 depicts the same events as observed for 21UR-1 in panel D.

### 5’-end processing of precursor-21UR-1 molecules follows a 3’-end processing step by ENDU-1

Depletion of XRN-2 resulted in decreased abundance of mature piRNAs with a concomitant accumulation of the corresponding 30-nt intermediate-precursors (**Figure 2A, 2B, 2E**), and strikingly without any change in the levels of the precursor-transcripts (**Figure 2A, 2B**), wherever we could detect them. This suggested that XRN-2 doesn’t influence the precursor transcription as well as its processing into the shorter intermediate (30-nt), rather works downstream of this processing activity. Since, these 30-nt intermediate-precursor bands were not smearish in appearance, rather by and large formed sharp bands on denaturing gels (**Figure 2A**), we hypothesized that an endoribonuclease activity is acting on the 60-nt precursor-transcripts to form sharp intermediate products. To identify this activity, we performed a candidate RNAi screen employing the aforementioned *C. elegans* 21UR-1/ piRNA reporter strain (**9**). We hypothesized that a highly extended 3’-end beyond the mature sequence due to the absence of such a processing activity might affect 21UR-1 functioning, unlike the PARN-1 perturbation situation, where the 21UR-1/ piRNAs get extended in their 3’-ends by only a few nucleotides. Amongst the candidate genes (**Supplementary Table 3**), depletion of ENDU-1 led to GFP desilencing in the germlines of a modest number of experimental worms [17% (±1.23, n = 50, N = 3)], although, desilenced signals were much lower in magnitude compared to that observed upon depletion of PRG-1 or XRN-2 (positive controls) (**Figure 4A**). This outcome suggested a possible role of ENDU-1 in piRNA metabolism, and indeed upon its depletion, the mature 21UR-1 accumulation was diminished with a simultaneous accumulation of the 60-nt precursor-transcript (**Figure 4B, 4C**). Surprisingly, unlike the control situation, the 30-nt intermediate-precursor couldn’t be detected in the experimental samples. We presumed that the reduced amount of 30-nt intermediate-precursors that got produced in the ENDU-1 depleted worms could readily undergo further processing by the wild type levels of XRN-2 and PARN-1 to produce the mature species. Depletion of ENDU-1 also resulted in diminishment of the mature 21UR-3372, a piRNA that is transcribed as an 85-nt long RNA species (**14, 16**), with a concomitant increased accumulation of its precursor-transcript (**Figure 4D, 4E**). Contrastingly, 21UR-1949 remained completely unaffected upon ENDU-1 depletion, which suggested that it plays a dedicated role towards processing of the precursor piRNAs that are born as longer transcripts (≥60-nt) (**Figure 4E**). Notably, it is possible that the modest depletion of mature 21UR-1 (**Figure 4B, 4C**) upon *endu-1*(*RNAi*) in reporter worms did not allow derepression of GFP to a higher magnitude and in a greater percentage of worms (**Figure 4A**). Here, long half-life of ENDU-1 could be the underlying reason, but we could not check its level due to the absence of an antibody.

**Figure 4.**
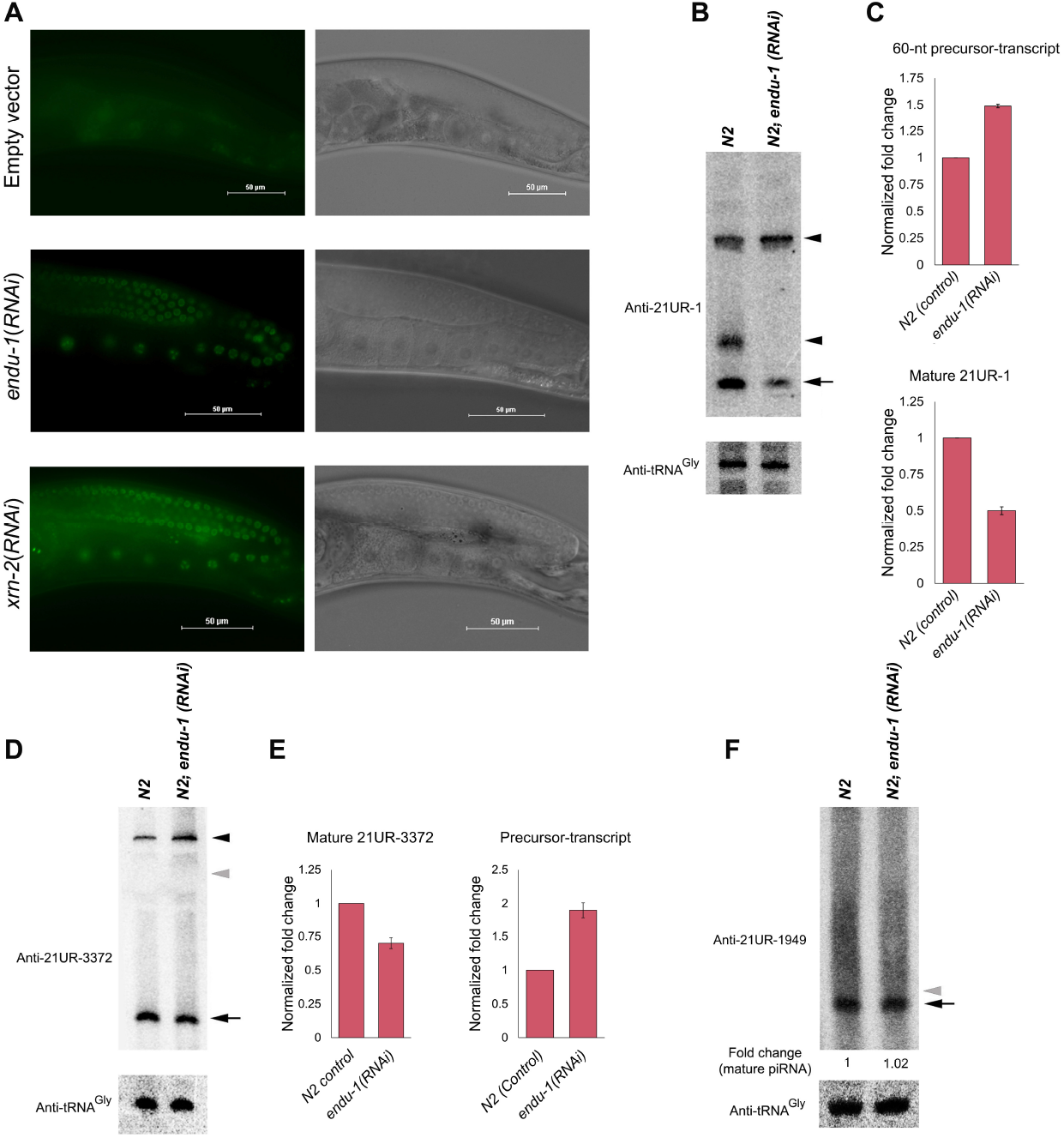
*endu-1*(*RNAi*) results in the depletion of the mature forms of those piRNAs that are born as long transcripts (≥60-nt) with a concomitant accumulation of their precursor-transcripts. **A.**Modest derepression of GFP-reporter gene (GFP-H2B) upon *endu-1*(*RNAi*). **B.**A representative northern blot showing depletion of mature 21UR-1 (arrow) and concomitant accumulation of the 60-nt precursor-transcript (top arrowhead) in worms undergoing *endu-1*(*RNAi*). The 30-nt intermediate-precursor (bottom arrowhead) is not detected in this experimental sample. tRNA^Gly^served as loading control. **C.**Graphical representations of the relative changes in the mature and precursor forms of 21UR-1 upon depletion of ENDU-1 (n = 3; means ± SEM). **D, E.**Observations similar to 21UR-1, as depicted in B and C, are made for 21UR-3372. The grey arrowhead in panel D indicates migration of a 65-nt long marker band. **F.***endu-1*(*RNAi*) has no effect on the mature form (arrow) of 21UR-1949 that is thought to be transcribed as a species much shorter than 60-nt. Grey arrowhead indicates migration of a 24-nt long marker RNA.

### Orchestration between XRN-2 and PRG-1 is critical for the generation of PRG-1 loaded incompletely processed precursors

It has been reported that PRG-1 is not involved in the transcription of 21U-RNAs, but is absolutely required for the stability of mature piRNAs (**15, 17**). In the absence of PRG-1, worms completely lack mature piRNAs (**3, 4, 15, 17**), and this could be due to the following two reasons: 1) PRG-1 facilitates precursor (eg. 30-nt intermediate-precursor of 21UR-1) processing, and in its absence they do not get processed into mature forms, and in turn get degraded. 2) It is also possible that the processing of precursor piRNA remains unaffected in the absence of PRG-1, but the processed mature piRNAs get degraded due to the absence of this effector protein.

To achieve clarity on the above-mentioned possibilities, we extracted total RNA from *prg-1*mutant worms and performed northern probing against 21UR-1. We observed no accumulation of mature 21UR-1, compared to wild-type control, when there was very little or no change in the level of the 60-nt precursor-transcripts (**Figure 5A, top panel, compare lanes 1 and 2**). Notably, these *prg-1* mutant worms grown at 20°C also did not accumulate any 30-nt intermediate-precursor, but when grown at an elevated non-permissive temperature (26°C), they displayed a substantial accumulation of this RNA species (**Figure 5A, top panel, compare lanes 2 and 4**). These results not only suggested that the initial 3’-end processing of the 60-nt precursor-transcript is independent of PRG-1, but also corroborated with the previous observation that PRG-1 is not involved in the 21U-RNA precursor transcription (**15**). These observations further suggested that PRG-1 might play a role in the processing of the 30-nt intermediate-precursors into 21-nt long mature forms. We hypothesized that after the initial 3’-end processing of the 60-nt precursor-transcripts, the 30-nt intermediate-precursors get loaded onto PRG-1, and the *bona fide* termini of the mature forms might remain protected by PRG-1. Thereafter, the unprotected nucleotides at the 5’-end along with the cap structure are likely to get processed out by XRN-2. And, in the absence of protection from PRG-1, XRN-2 holds the potential to degrade the entire 30-nt intermediate-precursor. To verify this hypothesis, we depleted XRN-2 in *prg-1* mutant background and probed for 21UR-1. Irrespective of the temperature at which the worms were grown, *prg-1; xrn-2(RNAi)*worms showed massive accumulation of the 30-nt intermediate-precursors, without any significant change in the levels of 60-nt precursor-transcripts (**Figure 5A, top panel lanes 3, 5; bottom panel**), which clearly indicated that in the absence of PRG-1 the 30-nt intermediate-precursors indeed get degraded by XRN-2 and this event is independent of the processing of the 60-nt precursor-transcripts. Here, it was clear that XRN-2 is capable of recognizing, directly or indirectly, the 30-nt intermediate-precursors in the absence of PRG-1. Thus, simultaneous loading of PRG-1 would be necessary to prevent the degradation of the substrate. However, if XRN-2 binds the 30-nt intermediate-precursor before PRG-1 does, in order to prevent immediate degradation, there has to be some involvement of ancillary factors. Furthermore, additional mechanisms, and possibly some more factors, would be required to subsequently allow XRN-2 to perform the required 5’-end processing of the precursor upon joining of PRG-1. Of note, it has been reported that the endoribonuclease activity of XRN-2 is dependent on substrate induced conformational changes (**24**), which might get affected at non-permissive temperatures, and that in turn might have caused the modest accumulation of the 30-nt intermediate-precursor in *prg-1* mutant worms grown at 26°C (**Figure 5A, top panel lane 4**).

**Figure 5.**
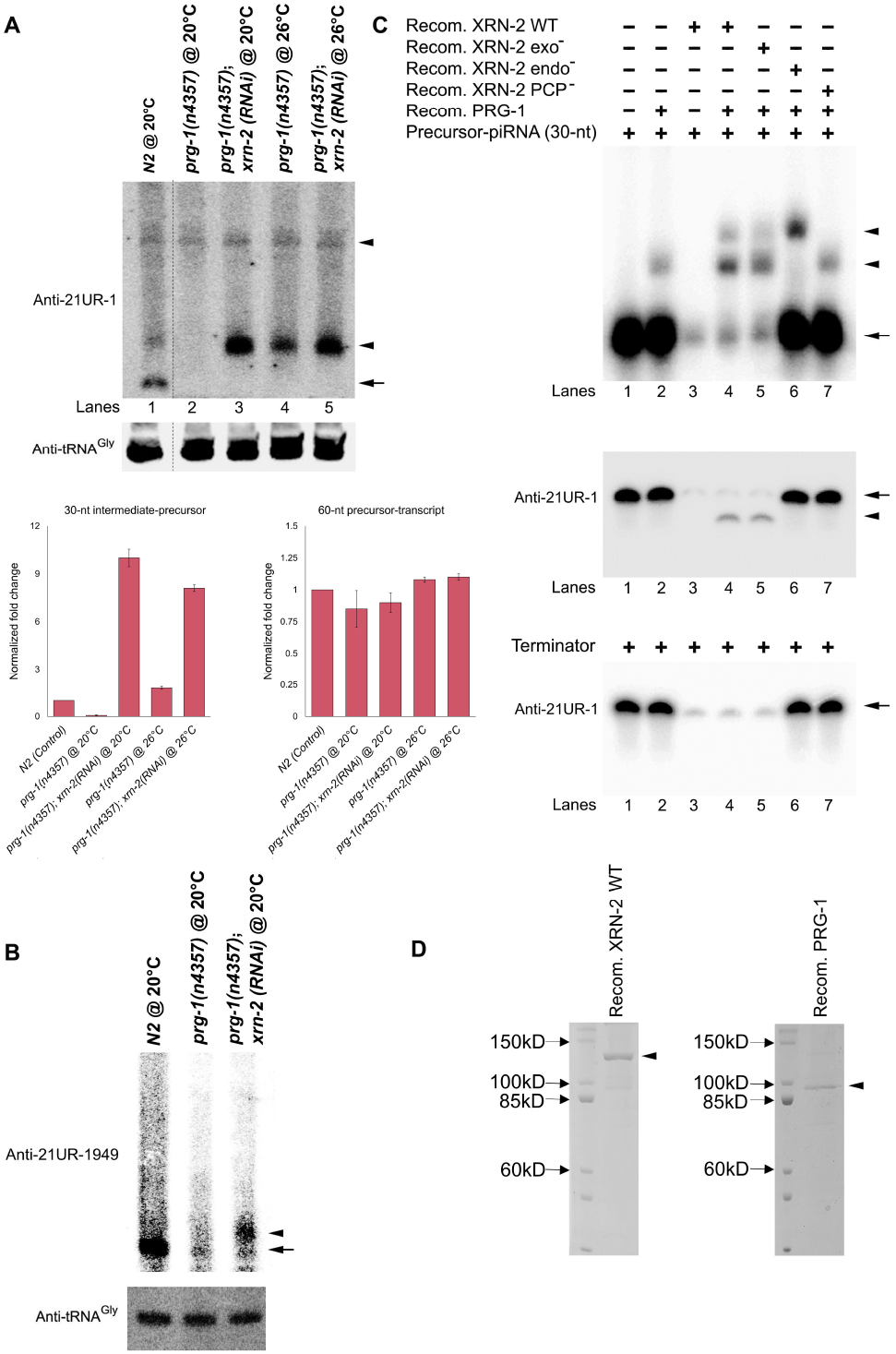
Orchestration between XRN-2 and PRG-1 is critical for the generation of PRG-1 loaded incompletely processed precursors. **A.**Accumulation of mature 21UR-1 depends on PRG-1, and it does not influence processing of the 60-nt precursor-transcript into 30-nt intermediate-precursor, whose accumulation is dependent on XRN-2. Anti-21UR-1 northern probing depicts absence of mature (arrow) and 30-nt precursor-transcript (bottom arrowhead) in *prg-1* mutant worms grown at 20°C (compare lanes 1 and 2). The 30-nt intermediate-precursor accumulates (lane 6), when the mutant worms are grown at 26°C, and a stronger accumulation is visible upon depletion of XRN-2 in the same genetic background (compare lanes 3,5 with lane 4). The 60-nt precursor-transcript (top arrowhead) levels remain largely comparable in all the samples. tRNA^Gly^served as loading control. All lanes are from the same blot. Bottom panel depicts graphical representation of these two precursor levels in the indicated samples (n = 3; means ± SEM). **B.**Mature 21UR-1949 is heavily depleted in *prg-1* mutant worms (middle lane), but a precursor form accumulates upon XRN-2 depletion in the same genetic background. These observations are in line with A. **C.Top panel.**5’-capped and radiolabeled 21UR-1 precursor RNA (30-nt) is incubated with proteins as indicated and the products are resolved on an EMSA gel. The substrate RNA is indicated with an arrow and the RNP bands with arrowheads. **Middle panel.**The same reactions, as indicated in the top panel, are performed with cold substrate RNA. After incubation, the RNA is extracted, resolved on a denaturing gel, transferred to a membrane, and subjected to northern probing for mature 21UR-1. **Bottom panel.**RNA extracted from the incubation reactions, as indicated in the middle panel, are treated with 5’-3’ Terminator nuclease, re-extracted, resolved on a denaturing gel, and subjected to northern probing as indicated. **D.**CBB stained recombinant wild type XRN-2 and PRG-1 employed in C.

An observation similar to that depicted for 21UR-1 in Figure 5A could also be made for 21UR-1949, which gets produced as a much shorter transcript (∼30-nt, **Figure 5B**). Thus, XRN-2 not only processes the 5’-ends of precursor piRNAs, but also functions as a part of a potential surveillance/ clearance mechanism, where they specifically degrade those ∼30-nt precursors that are inappropriately bound or not associated with the effector protein PRG-1, and thus contributes to piRISC homeostasis. Of note, upon northern probing, the sum of the signals for mature and the (∼30-nt) precursors observed for a given piRNA in *xrn-2*(*RNAi*) samples, always exceeded the sum of those signals observed in the control samples (**Figures 2A, 2E**). It is possible that a significant amount of the precursors undergo degradation by XRN-2 as a measure of quality control, and they also accumulate and contribute to the higher signal observed in *xrn-2*(*RNAi*) samples. Here, our observations reiterated the fact that not only the machinery towards transcription of the precursors, but the effective concentrations of all the RNA binding factors, processing enzymes and the effector protein PRG-1 indeed play critical roles in determining the abundance of mature piRNAs.

Our *in vivo* experiments also suggested that the 5’-end processing of the 30-nt intermediate-precursor by XRN-2 would be a prerequisite for PARN-1-mediated 3’-end trimming to form the mature 21-nt piRNA (21UR-1, **Figure 2C**), and under some given circumstances a complete 3’-5’ degradation of this precursor by the same enzyme may take place in the absence of the 5’-end processing (**Figure 3D, 3E**). It is very likely that in the wild type scenario, the PARN-1 is not able to work beyond its 3’-trimming activity due to occupation of the 3’-end of the precursor by PRG-1 that protects the nucleotides corresponding to the 3’-end of the mature piRNA. Importantly, the role of XRN-2 and its interaction with PRG-1 might determine whether the 3’-end of the precursor would be available for a limited or extensive action by PARN-1 (**Figures 3, 5A, 5B**). Therefore, to gain insights into the dynamics of interactions between the 30-nt intermediate-precursor, XRN-2 and PRG-1, we performed binding assays and resolved those reactions in native gel to reveal the different RNPs formed under different conditions (**Figure 5C, 5D**).

PRG-1 formed a weak RNP-band with the 30-nt intermediate-precursor. Whereas, in the presence of XRN-2, a stronger RNP-band was formed that co-migrated with the RNP formed in the presence of only PRG-1 (**Figure 5C, top panel, compare lanes 2, 4**). Additionally, a larger RNP-band did also get formed. We performed this reaction with cold substrate RNA and subjected the products (extracted RNA) to northern probing after resolving them under denaturing conditions, and we could detect the 30-nt intermediate-precursor as well as a shorter RNA, where the latter RNA species accumulated more than the former (**Figure 5C, middle panel lane 4**). Moreover, this shorter RNA was 5’-end processed, as it was sensitive to terminator nuclease treatment, whereas the 30-nt intermediate-precursor was resistant owing to its 5’-cap (**Figure 5C, bottom panel lane 4**). Notably, western probing revealed that the larger RNP-band housed both XRN-2 and PRG-1, whereas only PRG-1 could be detected in the smaller RNP-band (**data not shown**). These band patterns obtained on the native gel as well as through northern probing remained unchanged, when we replaced wild type XRN-2 protein with the one mutant for the exoribonuclease activity (**Figure 5C, compare lane 5 of top, middle, and bottom panels**), and that indicated to a role for the endoribonuclease activity of XRN-2. Interestingly, when we employed the endoribonuclease mutant XRN-2, we observed formation of the larger RNP-band only, housing both the proteins (**Figure 5C, top panel lane 6; data not shown**). Moreover, the northern probing of the reaction performed with cold 30-nt intermediate-precursor revealed the absence of the shorter processed transcript derived from the employed substrate (**Figure 5C, middle panel lane 6**). This essentially confirmed that the endoribonuclease activity of XRN-2 is required for the generation of the shorter RNA species devoid of the 5’-cap, and in the absence of this cleavage activity the RNP-band representing the ternary complex got stabilized. Contrastingly, in the presence of endoribonuclease activity (**Figure 5C, lanes 4, 5 of top and middle panels**), substantial accumulation of the binary complex comprising PRG-1 and mostly the 5’-processed shorter precursor RNA took place, where identity of the RNA could be confirmed upon gel extraction of the RNP band followed by analysis of the extracted RNA using denaturing PAGE (**data not shown**). It is possible that the energy released from the cleavage of the precursor-phosphodiester bond by the endoribonuclease activity of XRN-2 elicited a conformational change in the ternary complex that led way for the dissociation of XRN-2, and formation of a binary complex comprising PRG-1 and the 5’-processed shorter precursor RNA (**Figure 5C, compare lanes 4 and 6 in all panels**). Finally, when this binding reaction was also performed using PRG-1 and XRN-2 protein mutant for its RNA binding motif indispensable for its endoribonuclease activity, it resulted in the formation of a weak RNP-band that co-migrated with the band formed only in the presence of PRG-1, and only unprocessed 30-nt intermediate-precursor RNA could be detected (**Figure 5C. compare lanes 2 and 7 in all panels**). This observation suggested that XRN-2 directly binds RNA and facilitates binding by PRG-1. However, our observations do not reveal whether these proteins sequentially or simultaneously load on the RNA, or do they form an association in the first place and then load on the substrate RNA.

### XRN-2 depletion affects piRNA-dependent and independent endo-siRNAs, as well as genome integrity

Worms lacking active piRNA pathway are superficially normal but gradually become sterile over several generations as the germline loses its immortal nature (**3, 4, 9, 11, 12**). Contrastingly, depletion of XRN-2 through RNAi results in immediate sterility of the worms (P0) undergoing the treatment, which indicates that apart from contributing in piRNA biogenesis, it plays additional roles that might be critical for germline metabolism. It has been suggested that piRNAs have the potential to target/ interact with all the germline expressed transcripts (**5**), and it was observed that around two thousand gene-transcripts show decreased accumulation of endo-siRNAs (22G-RNAs) in *prg-1* mutants (**7, 8, 9, 13, 26**). Later on, ∼80% of the germline expressed transcripts (8986 out of 11,088) were suggested to be targeted and regulated by endo-siRNAs (**29, 30**). Thus, for obvious reasons, we examined the abundance of 22G-RNAs in XRN-2 depleted worms, at the same time point from where we already have reported the levels of their piRNAs (**Figure 2**). Compared to wild-type animals, in *xrn-2(RNAi)* worms we observed greater than 30% reduction in 22G-RNAs that might target as many as 4377 protein-coding genes. Notably, different AGO proteins bind to different subsets of 22G-RNAs and facilitate gene silencing or gene licensing, which might get affected by XRN-2 in a direct or indirect manner.

We first examined the status of PRG-1 dependent 22G-RNAs in XRN-2 depleted worms. Amongst the 4377 genes that might get affected by XRN-2, a significant overlap of 919 genes could be observed with the genes previously identified as PRG-1 dependent 22G-RNA targets (**Figure 6A, references 7, 9, 26**). We also examined the effect of XRN-2 depletion on a published list of 383 piRNA targets (**7, 9**), of which 256 showed a significant reduction in 22G-RNAs that could be mapped to them in *xrn-2(RNAi)* worms. This observation again supported that 5’-processing is indeed essential for the proper functioning of the piRNA pathway. In XRN-2 depleted samples, we indeed observed a widespread reduction in 22G-RNAs mapping to genes that are known to be targeted by the WAGO (WAGO-1/9) associated 22G-RNAs (1567 genes out of 2907 known targets, **Figure 6B, references 31, 32**).

**Figure 6.**
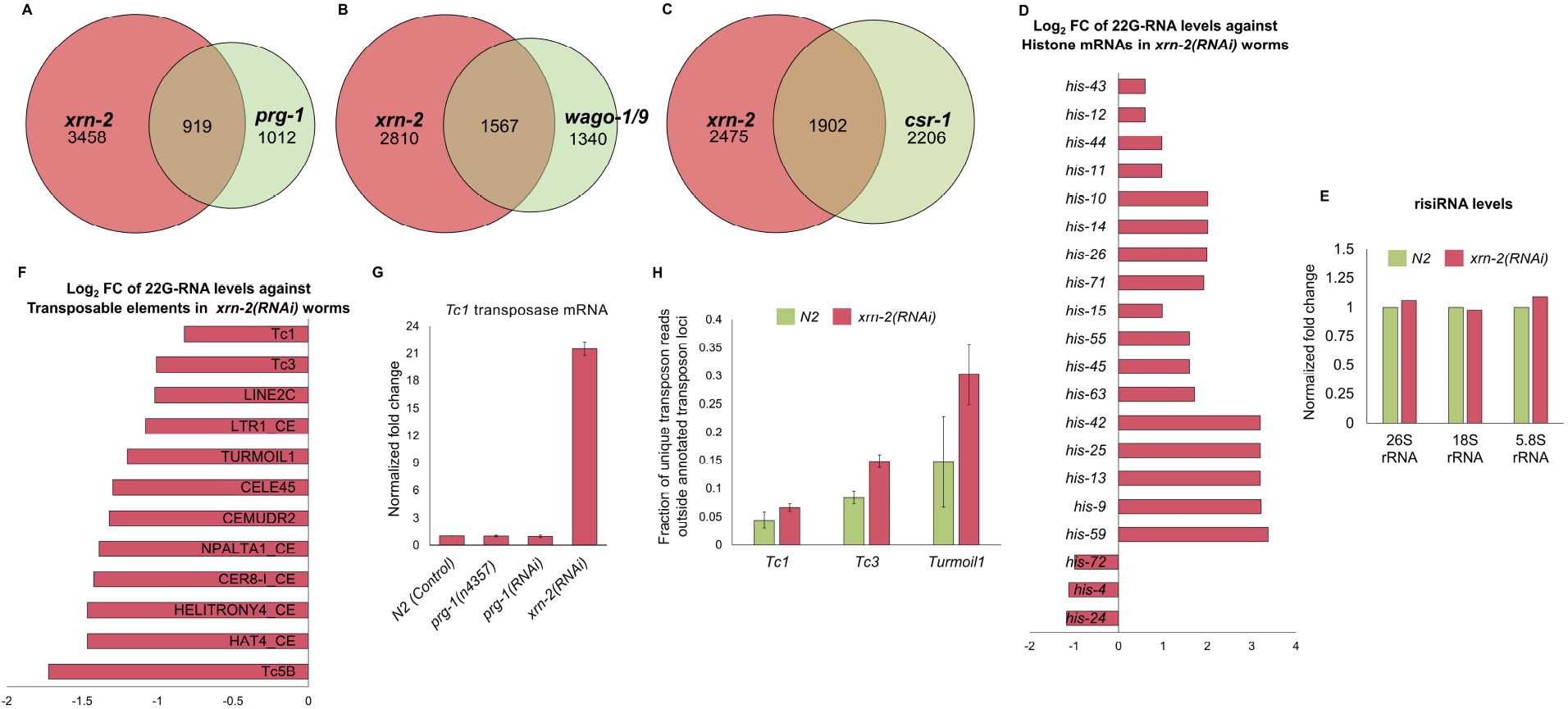
XRN-2 depletion affects piRNA-dependent and independent endo-siRNAs, as well as genome integrity. **A.**Venn diagram summarizing the overlap in the number of genes, whose 22G-RNA levels decrease in both *xrn-2(RNAi)* worms and *prg-1* mutant worms with respect to wild-type control. **B.**Venn diagram summarizing the overlap between genes targeted by WAGO-1/WAGO-9-associated and XRN-2 dependent 22G-RNAs. **C.**Venn diagram indicating overlap of genes depleted of 22G-RNAs in both CSR-1 mutant and XRN-2 depleted worms compared to wild-type control. **D.**Bar plot displaying log_2_fold change (log_2_FC) in 22G-RNA levels targeting different histone genes in XRN-2 depleted worms relative to wild type worms. **E.**Graphical representation of the normalized fold changes in the levels of risiRNAs against the indicated rRNAs upon XRN-2 depletion. **F.**Bar plot showing log_2_fold change in 22G-RNA levels targeting different transposon families (top 12) in XRN-2 depleted worms relative to wild type worms. **G.**RT-qPCR quantification (n = 3; means ± SEM) reveals that *Tc1* transposase mRNA gets dramatically upregulated in *xrn-2(RNAi)* worms as compared to control, whereas modest changes are noted in *prg-1(RNAi)*, and *prg-1* mutant worms with respect to control. **H.**Fraction of unique reads corresponding to *Tc1, Tc3* and *Turmoil1* transposons in non-transposon loci increases significantly in *xrn-2* (*RNAi*) worms as compared to control.

Surprisingly, our analyses also revealed that XRN-2 affects the CSR-1 associated 22G-RNAs that are produced independent of PRG-1 in a different pathway. Out of 4108 genes that get targeted by CSR-1, 1902 genes show reduced 22G-RNA levels in XRN-2 depleted worms (**Figure 6C, references 33**). Gene Ontology term enrichment analyses (GO term enrichment analyses) of these 1902 genes revealed that they are involved in various important cellular pathways including DNA replication and repair, RNA transport and surveillance, FoxO signaling pathway etc (**Supplementary Figure 4A, references 34**). CSR-1 has been proposed to oppose the engagement of piRISC complexes on transcripts and is thought to license genes for germline expression (**7, 8, 13**). Therefore, depletion of CSR-1 class of 22G-RNAs might lower the expression of germline enriched ‘self-transcripts’ essential for fertility. Whereas, reduction of WAGO associated 22G-RNAs may cause derepression of the transcripts meant to remain repressed. Of note, XRN-2 might be involved in the biogenesis of the CSR-1 associated 22G-RNAs or formation of active CSR-1-22G-RNA RNP complexes, and whose depletion results in the reduction and destabilization followed by degradation, respectively, of those effector RNA molecules. Overall, the deregulation of WAGO and CSR-1 pathways are likely to create grave imbalance in the transcriptomic landscape of the germline, and thus leading to sterility of the organism.

We observed an increase in 22G-RNAs mapping to a specific set of 1394 protein-coding transcripts in XRN-2 depleted samples, compared to control. Similar to the observations made in *prg-1* mutant worms, a majority of these 22G-RNAs were antisense to replicative histone mRNAs, which have been shown to be a major factor causing germline sterility (**Figure 6D; references 10, 26**). Additionally, GO term enrichment analyses indicated that the genes that were targeted by hyper-accumulated 22G-RNAs, beyond histones, are involved in axonal regeneration and Wnt signaling pathway (**Supplementary Figure 4B, reference 34**). Wnt signaling plays critical roles in multiple biological processes, including cell differentiation, migration, asymmetric cell division, neural development, organogenesis during embryonic development. Therefore, it is also possible that downregulation of the Wnt signaling pathway might have contributed to the sterility of the worms undergoing XRN-2 depletion. However, more work is necessary to understand which of these effects are linked to the deregulation of the endoribonuclease activity of XRN-2, and which ones are linked with its exoribonuclease activity. Importantly, we checked the level of 22G-RNAs that are antisense to rRNAs (also known as risiRNAs, **reference 12**), which remained largely unaltered in XRN-2 depleted worms (**Figure 6E**). This also corroborates with the previous observation that at our chosen time point for performing all the molecular analyses, depletion of XRN-2 exerted very little or no effect on the levels of mature 5.8s, 18s and 28s rRNA expression (**Supplementary Figure 1B**), and thus excludes the possibility that deregulated rRNA metabolism culminated in lack of progeny formation in the P0 worms undergoing *xrn-2* (*RNAi*).

We also found depletion in 22G-RNA levels in *xrn-2(RNAi)* sample against 1106 protein-coding genes, which are not known to be targeted by any of the well-studied Argonaute proteins. Employing GO term enrichment analyses (**34**), we noted that those genes are predominantly involved in phosphorus metabolism and several protein modification processes (**Supplementary Figure 4C**).

piRNAs and siRNAs are well known for their roles in silencing transposons (**1, 2**). However, in *C. elegans*, a handful of transposable elements (*Tc3, Turmoil1*) are thought to be regulated by 21U-RNA/ PRG-1 pathway (**3, 4, 9, 26**). To explore the role of XRN-2 in regulating the transposons, beyond the ones regulated by the piRNA pathway, we looked at the abundance of transposon-targeting 22G-RNAs in XRN-2 depleted worms. Compared to control animals, *xrn-2(RNAi)* worms showed substantial depletion of 22G-RNAs targeting as many as 79 transposable element families that are considered not to be regulated by the piRNA pathway (**Figure 6F**). Indeed, quantification using RT-qPCR revealed that XRN-2 depletion led to a dramatic accumulation of mRNA corresponding to the *Tc1* transposase (**Figure 6G**). This indicated to the possibility of higher expression and movement of the transposable elements in the experimental samples. To further check this, we also performed whole genome sequencing of control and *xrn-2(RNAi)* worms and found a significant increase in the fraction of unique transposon reads (corresponding to *Tc1, Tc3* and *Turmoil1*) outside annotated transposon loci in XRN-2 depleted worms as compared to control animals (**Figure 6H**).

## Discussion

Here, we have demonstrated the role of the endoribonuclease activity of XRN-2 in the biogenesis of piRNAs in *C. elegans*. Our study not only revealed how this activity contributes to the generation of the mature 5’-end of a piRNA, but also helped us to understand this step in the context of the entire maturation process, where it works in conjunction with other nucleases and the Piwi Argonaute protein PRG-1. The endoribonuclease activity of XRN-2 has already been implicated in rRNA and snoRNA metabolism in continuously growing worms, as well as microRNA metabolism during the dauer stage (**24, 25**). It is possible that XRN-2 protein housed in different complexes is participating in different RNA transaction pathways, where the other constituent subunits of the respective complexes might be performing distinct roles. Thus, deeper insights can be achieved, if the different steps of piRNA biogenesis would be studied from the context of specificity, functionality and regulatability of the concerned enzymes residing in their *bona fide* complexes.

Recently, a four-subunit protein complex (PETISCO/ PICS) has been shown to be important for the 5’-end processing of piRNA precursors in worms (**19-21**). The subunits of this complex lack any known decapping or nuclease activity required for the removal of a cap structure and precursor-sequence. Interestingly, a subunit (IFE-3) is known to possess cap-binding activity and was shown to bind the capped piRNA precursors. It is possible that PETISCO/ PICS first binds capped precursors and then XRN-2 gets recruited to process their 5’-ends. Alternatively, they work independently in parallel pathways. Intriguingly, we observed that some of the XRN-2 substrates (eg. 21UR-1949) indeed overlap with the precursors that are bound by IFE-3, which indicates that XRN-2 might be working in association with PETISCO/ PICS to process certain precursor molecules, whereas for other substrates (eg. 21UR-1), it is working independently.

We observed a large number of Type I/ Ruby-motif piRNAs (4475 out of 7439 detected) to be substrates of XRN-2, whose mature forms got depleted upon XRN-2 knockdown. 5’-unprocessed precursors of some of the affected mature piRNAs got accumulated at the same time. Notably, these precursors were underrepresented in our sequencing data, which could be due to their inherent instability or because of some cellular surveillance machinery actively removing unprocessed precursors or due to possession of some chemical modifications that excluded those precursors from getting cloned into the library. These XRN-2-substrate Type I/Ruby-motif piRNAs are uniformly distributed over the two defined loci in chromosome IV that harbor ∼15,000 piRNA genes. Currently, it is not clear that how only these precursors get processed by XRN-2, and not all. It is possible that there are parallel processing pathways and they might recognize their substrates due to the presence of some sequence motifs, though we were unable to confidently observe any significantly enriched motifs in the XRN2-targeted piRNAs from our sequencing dataset. Depletion of XRN-2 also led to the depletion of Type II/ non-Ruby-motif piRNAs (78 out of 133 detected species) that are produced from loci dispersed all over the genome. Many of these piRNAs overlapped with known TSSA RNA sequences (**data not shown**). This observation again suggested that the capped TSSA RNAs are indeed potential Type II/ non-Ruby-motif piRNA precursors if they would have an appropriately positioned U-residue to become the 1^st^ nucleotide of the mature 21U RNA/ piRNA (**14**), and appropriate machinery for the recruitment of the necessary biogenesis factors to these transcripts.

There is not enough clarity about the sizes of the precursor transcripts of piRNAs. High-throughput sequencing-based approaches detected relatively smaller ∼26-28 nt long piRNA precursors (**14, 15**). A recent study aimed to understand the role of the Integrator Complex in transcription termination of piRNA precursor-transcripts reported two distinct populations of piRNA-precursors (28-nt and 48-nt long, **reference 18**). Earlier, RACE-based methods had detected the presence of a ∼75-nt long precursor (pre-21UR-3372, **reference 16**). Here, employing northern probing, we could detect precursors in two size ranges: ≥60-nt long (eg. 21UR-1) and ∼30-nt long (eg. 21U-1949). The longer precursors (≥60-nt long) require an additional 3’-processing step that was previously unknown. Whereas, all the substrates, irrespective of their lengths, are dependent on the endoribonuclease activity of XRN-2 for the generation of their mature 5’-ends. To understand the mechanistic details of the processing events, we focused largely on 21UR-1 as a model piRNA and employing a combination of *in vivo* and *in vitro* experiments, we could demonstrate how its 60-nt long precursor-transcript is first acted upon by ENDU-1 to remove a long stretch of nucleotides at its 3’-end (**Figure 4**), and the resulting ∼30-nt long intermediate product is held up by XRN-2 and PRG-1, sequentially or simultaneously (**Figure 5**). XRN-2, owing to its endoribonuclease activity, removes the 5’-cap and the following two nucleotides to form the mature 5’-end, beginning with a Uridine that probably remains bound by PRG-1 (**Figures 2, 3, 5**). The 5’-processing event is followed by the PARN-1-mediated removal of the nucleotides in the 3’-end of the precursor molecule that are probably not bound and protected by PRG-1 (**Figures 2, 7; reference 17**). Interaction between these players might be transient, where energy released from the cleavage of the phosphodiester bonds might elicit critical conformational changes required for the formation of the RNP comprising PRG-1 and mature 21-nt long piRNA. It is very likely that additional protein factors and mechanisms are involved in the formation of the functional piRISC, whose revelation would be critical to understand piRNA metabolism.

**Figure 7.**
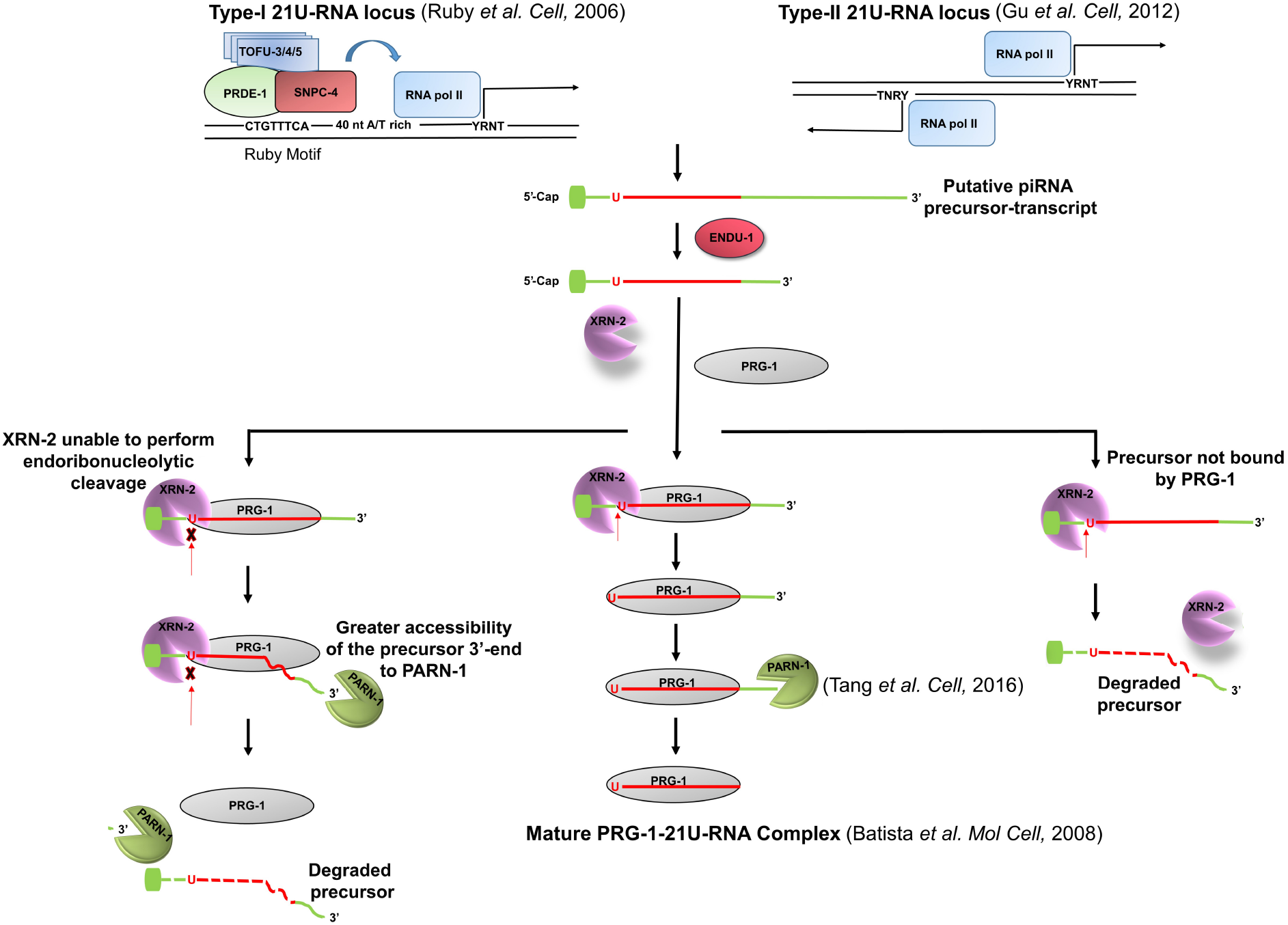
piRNA biogenesis model depicting formation of mature piRNA from a precursor-piRNA molecule (centre), as well as avenues for surveillance and clearance of under-processed and inappropriate (PRG-1 unbound) precursors (two sides). The scheme (**centre**) encapsulates the sequential processing of the piRNAs born as longer transcripts (eg. 21UR-1) by ENDU-1 (this study), XRN-2 (this study), and PARN-1 (**17**). ENDU-1-processed precursor undergoes further processing by XRN-2 and PARN-1 in association with PRG-1 to generate the PRG-1 bound mature form. Figure 5A suggests that XRN-2 has the potential to act on the relevant precursor in the absence of PRG-1 (**right side**), and 5C suggests that XRN-2 may facilitate binding of that same precursor molecules by PRG-1. Additional mechanisms and factors might be involved in coordination of these events, and precursor binding by both XRN-2 and PRG-1 at the same stage has been depicted for simplicity. Precursor molecule bound with XRN-2 that is incapable (indicated by cross-mark) of performing endoribonucleolytic cleavage (site indicated by red arrow) attracts PARN-1-mediated 3’-5’ degradation (**left side**). piRNAs born as shorter transcripts are independent of ENDU-1 (**Figure 4**), and might get acted upon directly by XRN-2, PRG-1, and PARN-1.

Similar to *prg-1* mutant worms (**3, 9, 11**), worms mutant for the endoribonuclease activity of XRN-2 [*xrn-2* (*PHX25*)] show temperature dependent mortal germline (*mrt*) phenotype (**24**). Whereas, XRN-2 depletion causes immediate sterility, irrespective of the growth temperature. Depletion of XRN-2 affects not only the abundance of mature piRNAs and piRNA-dependent endo-siRNAs, but piRNA independent endo-siRNAs as well. It is possible that the exoribonuclease activity of XRN-2 might have some role in the metabolism of piRNA-independent endo-siRNAs, and thus detailed analyses of the relevant exo/ endoribonuclease mutants would be an absolute requirement to understand their relative contributions. Interestingly, *xrn-2* (*RNAi*) worms show higher expression and movement of not only piRNA-regulated transposable elements, but also those that are known to be unlinked to piRNAs. It would be interesting to know how the expression and mobilization of the latter category of transposable elements are affected by XRN-2, and whether those jumping elements have any preferred target sequence motifs, which in turn might make certain gene families more vulnerable than others. Therefore, XRN-2 not only plays a critical role in shaping the germline transcriptome by modulating small RNA pathways, but it also contributes to the integrity of the genome by regulating the homeostasis of transposable elements.

## Author Contributions

SC designed the research. Experiments were designed by SC and AS, and performed by AS and TC. Together, all the authors analyzed the results and SC wrote the manuscript.

## Acknowledgments

This work was initiated with funds from DBT (Govt. of India) - IISc Partnership Program, then continued and sustained with personal funds. AS and TC received fellowships from CSIR (Govt. of India) and KVPY-MHRD (Govt. of India), respectively. Some strains were provided by the CGC, which is funded by NIH Office of Research Infrastructure Programs (P40 OD010440).

