## Supplementary Information for "Biogenesis of mature piRNAs in *Caenorhabditis elegans* is dependent on orchestration between multiple ribonucleases and PRG-1"

**Laboratory of RNA Molecular Biology**

**Indian Institute of Science**

**Bangalore, India**

**This file includes:**

Supplementary Figures 1 to 4

Materials and Methods

Supplementary Tables 1 to 5

**Supplementary Figures**

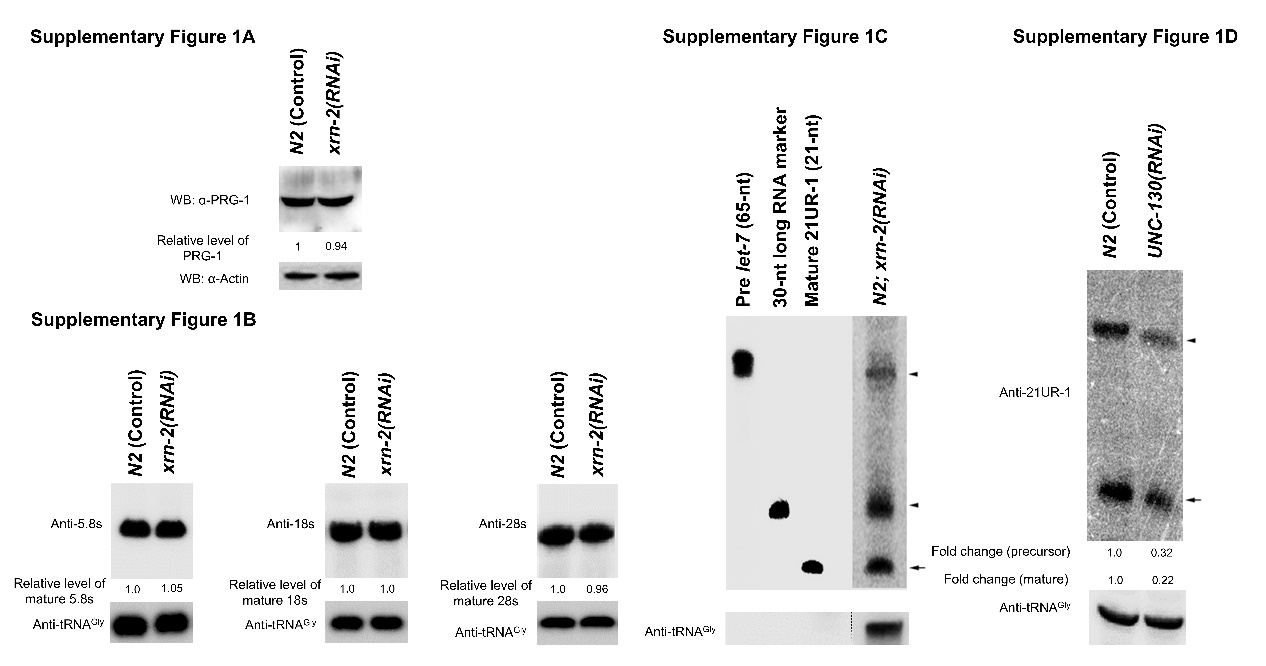

**Supplementary Figure 1**

**A.** *xrn-2*(*RNAi*) does not affect the expression of PRG-1 protein.

**B.** Northern blot analyses reveal no significant change in the levels of mature 5.8s, 18s and 28s rRNAs in *xrn-2(RNAi)* worms as compared to control worms, at the given time point. The indicated samples were resolved on a 1.5% Formaldehyde Agarose denaturing gel and subjected to northern probing. tRNA^Gly^ served as loading control.

**C.** Length estimation of 21UR-1 precursors. Total RNA extracted from *xrn-2(RNAi)* worms was resolved alongside radiolabeled synthetic pre *let-7* (65-nt), 30-nt long RNA marker and synthetic 21UR-1(21-nt), transferred to a membrane and only the relevant lane was probed against 21UR-1. The blot-pieces were realigned and subjected to phosphorimaging.

**D.** A representative northern blot showing depletion of both mature 21UR-1 (arrow) and 60-nt precursor-transcript (top arrowhead) in worms undergoing *unc-130*(*RNAi*), a known transcription factor involved in the transcription of piRNAs.

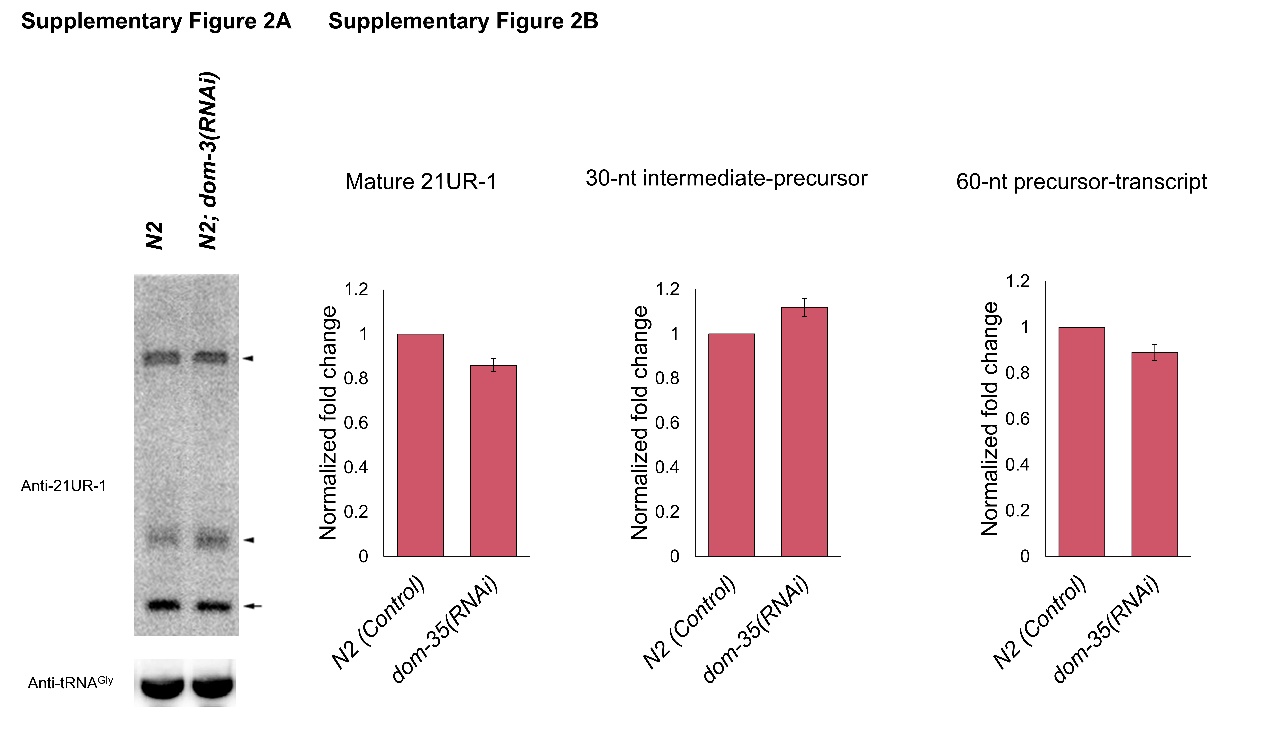

**Supplementary Figure 2**

**A.** A representative northern image depicting modest downregulation of mature 21UR-1 (arrow), modest increase in the level of the 30-nt intermediate-precursor (bottom arrowhead), and nearly unchanged level of 60-nt precursor-transcript (top arrowhead) in *dom-3*(*RNAi*) samples, compared to control (*N2* worms). tRNA^Gly^ served as loading control.

**B.** Graphical representation of the relative levels of the mature and precursor forms of 21UR-1 upon depletion of DOM-3 (n = 3; means ± SEM).

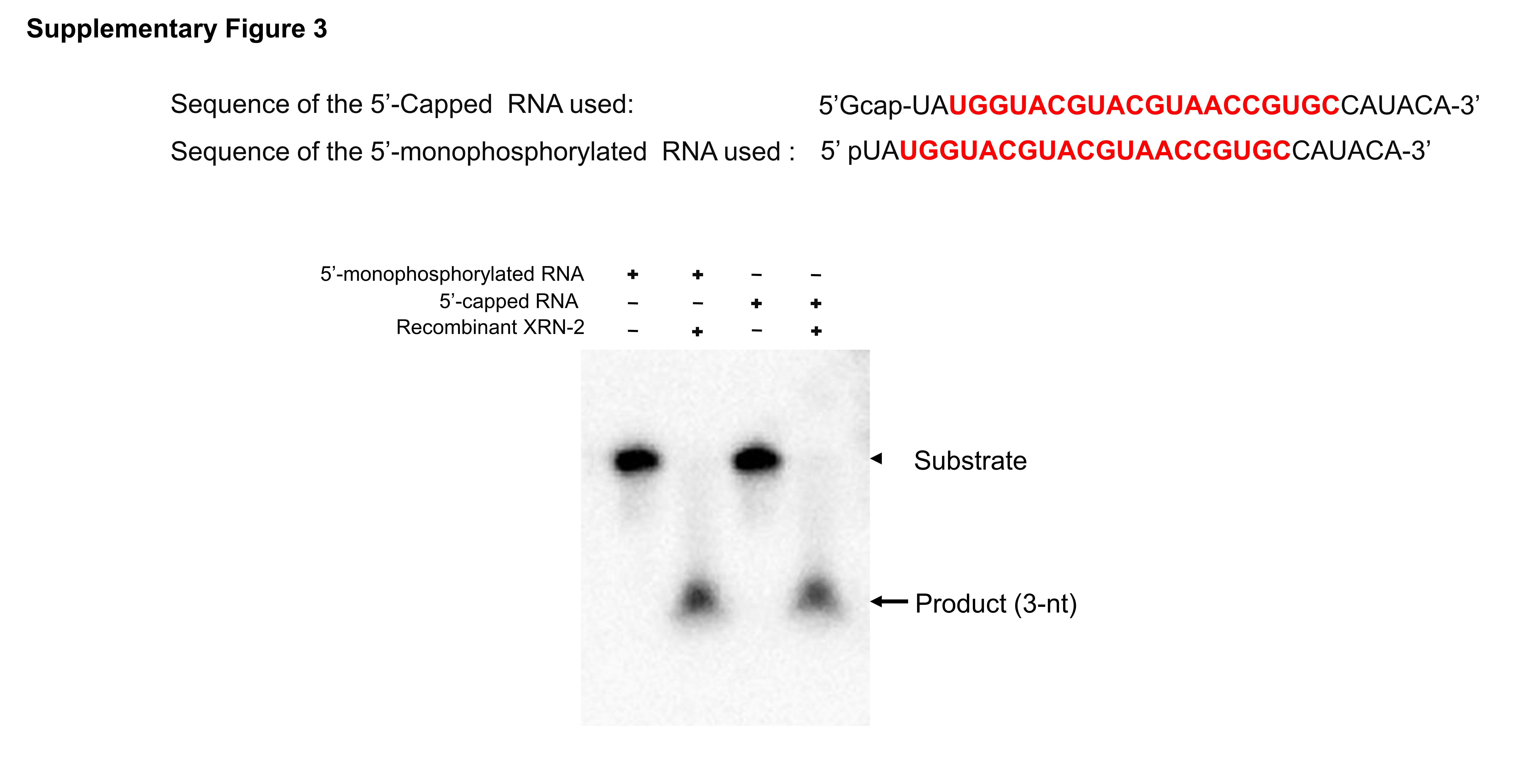

**Supplementary Figure 3**

Activity of recombinant XRN-2 on 5’-monophosphorylated and 5’-capped RNA substrates. Presence of 5’-cap does not affect the endoribonuclease activity of XRN-2.

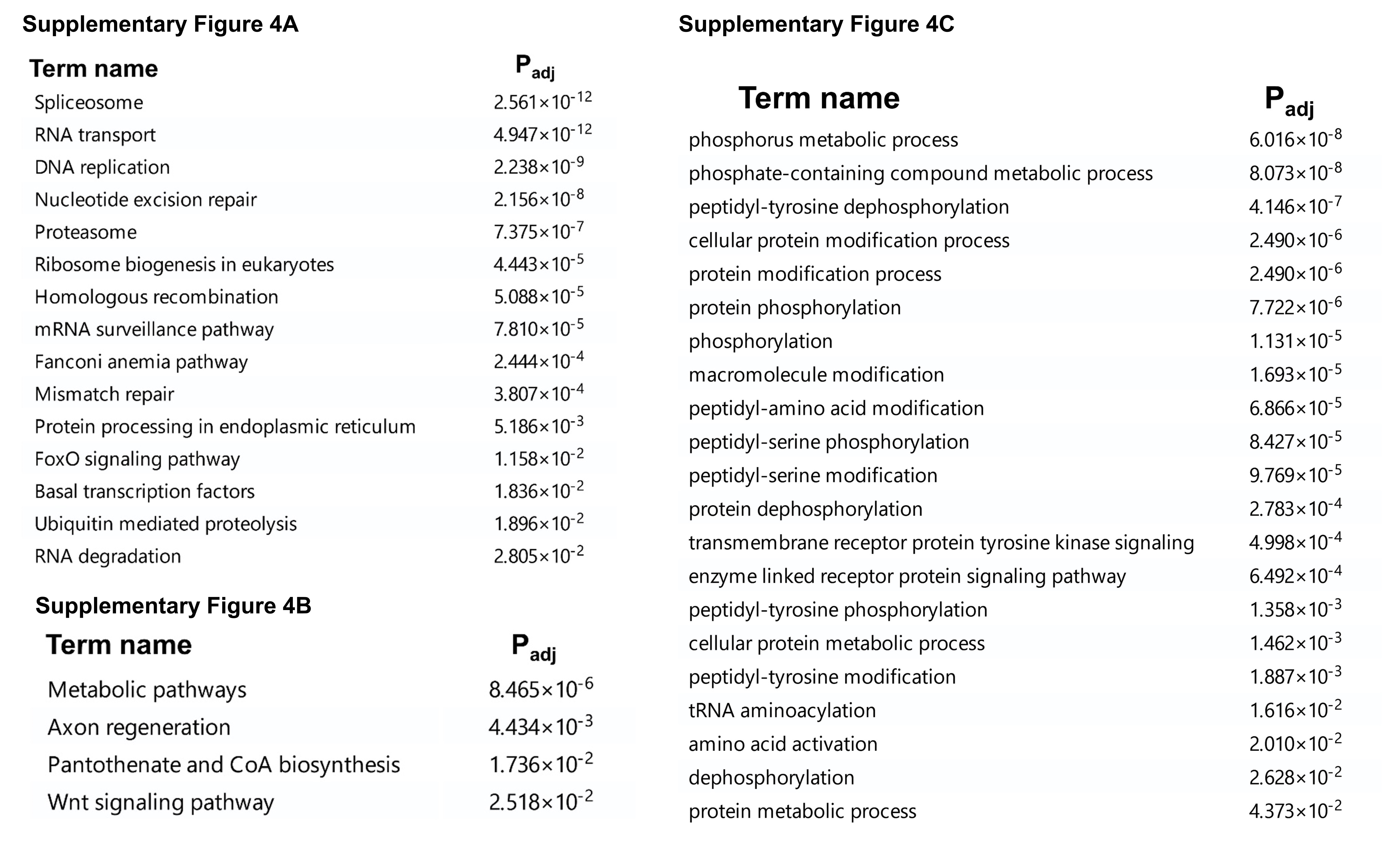

**Supplementary Figure 4**

**A.** The most significantly enriched KEGG pathways of 1902 genes affected by both CSR-1 and XRN-2.

**B.** The most significantly enriched KEGG pathways of 1394 genes that show upregulated 22G-RNA levels upon XRN-2 depletion.

**C.** Top 21 enriched biological functions of the 1106 genes that show downregulated 22G-RNA levels upon XRN-2 depletion, where the effector Argonaute proteins are currently unknown.

**Materials and Methods**

**Nematode Strains and Maintenance.** *C. elegans* were grown under standard conditions at 20°C using *Escherichia coli* strain OP50 as food source (**34**). Bleaching followed by starvation-induced L1 arrest was used to generate synchronized cultures. The worm strains used are described in **Supplementary Table 4.**

**RNAi experiments.** RNAi performed by feeding worms with *E. coli* strain HT115 (DE3) transformed with the control (L4440 vector) or L4440 vector construct containing specific gene sequences. RNAi clones were either used from GE Healthcare/ Dharmacon *C. elegans* RNAi Collection or were generated using L4440 vector and transformed in HT115 (DE3). Bacteria were grown overnight in LB with antibiotics at 37°C and seeded onto NGM plates containing 1.0 mM isopropyl b-d-thiogalactoside (IPTG), 100 mg/ml carbenicillin and 15 mg/ml tetracycline and incubated for 24 hours at room temperature. L1 larvae were placed on RNAi plates and grown at 20°C, if not otherwise indicated. For Northern blotting and RT-qPCR, worms were harvested at young-adult stage (50-52 hours). For candidate gene-RNAi screen, worms were grown on RNAi plates at 20°C up to adult stage and scored for germline associated *gfp* signal.

**RNA isolation, Northern blotting and RT-qPCR analysis.** For total RNA isolation, animals were harvested at young-adult stage from plates by washing with M9 buffer (22 mM KH_2_PO_4_, 22 mM Na_2_HPO_4_, 85 mM NaCl, 1mM MgSO_4_). Animals were pelleted and suspended in ten pellet volumes of TRIzol reagent (Ambion), and total RNA was extracted according to the manufacturer’s protocol. Northern blotting of endogenous RNA was done as described (**35**). In short, for detecting small RNA, ~50 μg of total RNA or 15-20 μg of mirVana (Ambion) enriched small RNA was resolved on 7 M urea/ 12% PAGE, transferred to Hybond N+ (GE Healthcare) membrane and probed appropriately. For detecting Ribosomal RNAs, 5.0 μg of total RNA was resolved using 1.5% Formaldehyde Agarose denaturing gel and transferred to Hybond N+ (GE Healthcare) membrane. Hybridization was carried out overnight using 5’-end labeled DNA probe.

For RT-qPCR, 2.0 μg of total RNA of the respective samples were reverse transcribed using SuperScript™ III Reverse Transcriptase (Invitrogen) in 20 μl reaction containing 5 mM DTT, 0.5 mM dNTPs, 2.5 μM oligo (dT)_20_ and 1.0 μl of enzyme, in accordance with the manufacturer’s protocol. 1.5 μl of each of the reverse-transcription reactions were amplified with PowerUp^TM^ SYBR Green Master Mix (Applied Biosystems) in a total volume of 10 μl using specific forward and reverse primers at a concentration of 0.5 μM each, and analyzed in a ViiA™ 7 Real-Time PCR System using ΔΔCt method. Values were normalized against the value of *act-1* expression. The experimental values, representing means (±SEM) from three independent biological replicates, were compared to the expression levels of the corresponding control samples. The primers used are described in **Supplementary Table 5**.

**Cloning and expression of recombinant XRN-2 and PRG-1.** cDNA was generated from *C. elegans* total RNA using SuperScript™ III Reverse Transcriptase (Invitrogen) and oligo (dT)_20_, and then PRG-1 was PCR amplified using gene specific primers and Q5 High-Fidelity DNA Polymerase (NEB), and cloned in pFastBac^TM^HT B (Invitrogen). The sequence confirmed correct ORF was further used to generate the recombinant bacmid DNA as per the manufacturer’s protocol (Bac-to-Bac Baculovirus Expression System, Invitrogen) towards expression of the recombinant N-terminal Strep tag containing PRG-1 proteins. Strep tagged PRG-1 and XRN-2 (wild type and mutants) were purified using Strep-Tactin^R^XT Superflow^R^ high capacity resin (IBA, Germany) and methods as described before (**24**). The primers used are described in **Supplementary Table 5**.

**Preparation of RNA substrates.** 30-mer precursor transcripts corresponding to 21UR-1 were transcribed from DNA cassettes using a MEGAshortscriptTM T7 transcription kit (Ambion) in presence of α-32P-UTP, cap analog m7G(5’)ppp(5’)G (Ambion) as per the supplier’s protocol. The DNA cassettes were prepared by annealing appropriate forward and reverse primers (The primers used are described below) followed by fill-in reactions using Klenow (NEB). Double stranded DNAs of appropriate length were then gel purified, PCR amplified, and cloned into TOPO TA vector (Invitrogen) and sequence confirmed. For *in vitro transcription*, the DNA cassettes were PCR amplified from the relevant TOPO TA vectors, gel eluted, and directly used as template.

**Sf-9 Cell culture.** Sf-9 cells were grown in Grace’s insect cell media (Gibco, Invitrogen) supplemented with 10% heat- inactivated FBS at 27°C in humidified atmosphere.

**Microscopy.** DIC images and fluorescent images were captured using Nikon NiE microscope and NIS AR software (Japan).

**Preparation of worm lysate.** Young adult worms grown on NGM plates seeded with OP50 *E. coli* strain were harvested and washed twice with M9 buffer. The worm pellets were then suspended in lysis buffer (50 mM Tris-Cl pH 8.0, 3 mM DTT, 3.0 mM MgCl_2_, 0.1% Triton X-100, 50 mM KCl, 5.0% glycerol, Protease inhibitor cocktail from Sigma and Roche) and ground in liquid nitrogen. After thawing, the samples were centrifuged at 30,000 x g for 20 min; the clear supernatants were collected and used for further experiments.

**Antibodies and Western blotting.** Bacterially expressed full length PRG-1 protein was injected in rabbit (New Zealand White Rabbits) as per the methods described before (**36**). The collected sera were processed as described before (**25**). For western blotting, α-PRG-1 serum was used at 1:500 dilutions, followed by the use of α-Rabbit HRP-conjugated secondary antibody (GE Healthcare) at a dilution of 1:10,000. Detection was performed with Amersham^TM^ ECL^TM^ Prime Western Blotting Detection Reagent, and ImageQuant LAS 4000 Chemiluminescence Imager (GE Healthcare).

**Specificity of PRG-1 antiserum.** Western blot performed using wild-type and *prg-1* mutant worm lysates, employing an antibody generated against PRG-1. In wild-type lysate, single band of desired size ( ̴ 93 kD) could be detected that was absent in the *prg-1* mutant worm lysate. Actin served as loading control.

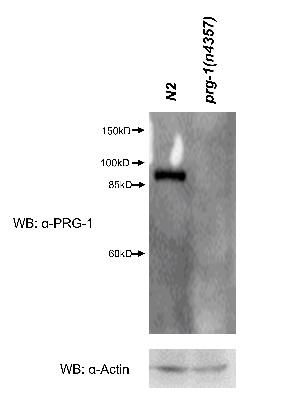

**Terminator experiment.** 50 μg of mirVana (Ambion) enriched small RNA samples were resolved on a 15% polyacrylamide/ 7M urea gel and 15–90 nt long RNA species were gel eluted. These gel purified species were incubated with Terminator^TM^ 5’-phosphate-dependent exonuclease (Lucigen) at 30°C for 1 hour, and then those reaction products were further resolved using denaturing gel, transferred to membranes, and probed with appropriate oligos.

***In vitro* binding/ processing assay.** Molar excess of *in vitro* transcribed body radiolabeled and 5’-capped 30 nt long RNA substrate was incubated with 50-75 μM of recombinant protein(s) at 4°C for 30-45 mins. Equimolar amount of recombinant proteins were used in reactions involving multiple proteins. The reactions were resolved employing 6% EMSA gel at 4°C in the presence of 5.0 mM MgCl_2_. For specific assays reactions were carried out with cold substrate and the reactions were splitted into two-halves. One part was resolved using denaturing UREA PAGE and probed for 21UR-1. The other part was subjected to treatment with Terminator^TM^ 5’-phosphate-dependent exonuclease (Lucigen) at 30°C for 1 hour, and then the products were further resolved using denaturing UREA PAGE and probed for 21UR-1.

**Small RNA Cloning.** Total RNA was extracted from young adult control and *xrn-2(RNAi)* using TRIzol reagent (Ambion). Small RNA was enriched using MirVana Kit (Ambion). RNAs ranging from 18–40 nt were further purified employing 7 M urea/ 15% PAGE. One μg each of purified small RNA samples were treated with RNA 5’- Pyrophosphohydrolase (NEB), which allows for cloning of 5’-capped/ triphosphate carrying small RNAs, at 37°C for one hour (**19**). The reactions were stopped with the addition of phenol:chloroform, and the treated small RNAs were purified via ethanol precipitation. Sequencing libraries were prepared using the NEBNext Multiplex Small RNA Library Prep Set for Illumina (NEB). PCR amplicons were size selected on 10% polyacrylamide gels. The libraries were sequenced on Illumina HiSeq platform.

**Genomic DNA extraction and sequencing.** The worm pallets were suspended in 500 μl lysis buffer [100 mM Tris-Cl (pH 8.0), 50 mM EDTA, 100 mM NaCl and 1% SDS], followed by 30 μl of 30 mg/ml proteinase K (Sigma), was added. The samples were incubated for 1.0 hour at 60°C until the tissue was totally dissolved. The DNA was then extracted from each sample with an equal volume of chloroform, followed by equal volume of phenol: chloroform: isoamyl alcohol solution (25:24:1). The upper aqueous layer was transferred to a fresh, sterilized microcentrifuge tube. RNase A (10 μl of 10 mg/ml; Sigma) was added, and the solution was incubated at 37°C for 1.0 hour. Finally, DNA was isolated by phenol/chloroform and chloroform extractions, followed by ethanol precipitation in the presence of 0.3 M Sodium Acetate. Sequencing libraries were prepared using the QIAseq FX DNA Library Kit (QIAGEN). The amplified libraries were purified using JetSeq Beads (Bio, # 68031). Further, the Illumina-compatible sequencing libraries were quantified by Qubit fluorometer (Thermo Fisher Scientific, MA, USA) and the fragment size distribution was analyzed on Agilent TapeStation 2200. The libraries were sequenced on Illumina HiSeq platform.

**Small RNA-Seq data analysis.** Reads were adapter-trimmed using fastp v0.20.0 (fastp -w 8 -i ${sample}.fq.gz -o ${sample}.trim.fq.gz --length_required 10) [2] using automatic adapter detection. Reads smaller than 10 nt after adapter and quality trimming were discarded. Remaining reads were aligned to *C. elegans* genome release WS280 using bowtie v1.2.3 (bowtie -p 8 -v 1 --tryhard --all --best --strata -S --no-unal ./index/ref_genome_ws280 ${sample}.trim.fq.gz) [3] allowing maximum of 1 mismatch in the complete alignment. Read alignments were then processed by a custom Python script relying on pysam 0.16.0.1 [4] to count alignments corresponding to known miRNA and piRNA genomic loci from release WS280 as well as the loci described in Gu et al, 2012 [5] (after converting the coordinates from Wormbase release WS215 to WS280 using the Wormbase remap-between-versions script). piRNA reads which mapped starting at 1 or 2 nt upstream of the annotated 21U locus were considered precursor reads, and reads mapping from the start of the annotated locus up to 5 nt downstream were considered as mature reads (3’ extended piRNAs remain functional [6]). siRNA reads were also obtained from these read alignments by considering all aligned reads which did not map to known miRNA or piRNA loci, possessed a leading ‘G’, were between 20 to 23 (22G) and 24 to 27 (26G) in length, and mapped antisense to an annotated genomic locus. As multiple alignments were considered, counts were normalized against total reported alignments (either miRNA + piRNA or siRNA) and the read count of an external spike-in hsa-miR-122 and expressed as RPM. Fold changes were computed directly from RPM due to having a single set of data.

**Whole Genome Sequencing data analysis.** Reads were adapter-trimmed using fastp v0.20.0 (fastp --in1 "${sample}_R1.fq.gz" --in2 "${sample}_R2.fq.gz" --stdout --dont_eval_duplication --correction --thread 8) [2] using automatic paired-end adapter detection and base correction. The processed reads were aligned to *C. elegans* genome release WS280 using bowtie2 v2.3.5.1 (bowtie2 -x ./index/ref_genome_ws280 --interleaved - --local -p 8) [3] using the --local preset. PCR and optical duplicates were marked (and removed) using the samtools duplication pipeline (samtools collate -f -@ 8 -O - | samtools fixmate -rcm -@ 8 - - | samtools sort -l 9 -@ 8 | samtools markdup -l 150 -d 100 -rS -@ 8 - - > "${sample}.bam"). The remaining aligned reads were then analysed by a custom Python program to discover alignments containing transposon sequences. Specifically, reads possessing greater than 70% overlap with 150 nt of the 5’ or 3’ sequences of Tc1, Tc3, and Turmoil1 were considered to possess transposon insertions and their alignment coordinates to the genome were recorded and mapped to annotated transposon and other genomic loci as per Laricchia et al, 2017 (after converting the coordinates from WS245 to WS280 using the Wormbase remap-between-versions script) and WS280 respectively in a non-strand-specific manner. The transposon mobilization was calculated as a fraction of transposon flank-containing reads mapped to non-transposon (of the same transposon family as the flank) loci versus the total number of transposon-flank containing reads detected. Only the first alignment of each aligned read was processed to make downstream analysis simpler – bowtie2 reports the best alignment by default.

Candidate genes employed in piRNA derepression screen (Related to Figure 1).

**Supplementary Table 1**

| **Name** | **Mean** | **SEM** |
| --- | --- | --- |
| *xrn-2* (5'-3' exoribonuclease) | 66.00% | 4.11% |
| *xrn-1* (5'-3' exoribonuclease) | 2.00% | 0.71% |
| *dom-3* (Decapping enzyme) | 18.46% | 0.63% |
| *eol-1* (Decapping enzyme) | 0.00% | 0.00% |
| *dcap-1* (Decapping enzyme) | 0.00% | 0.00% |
| *dcap-2* (Decapping enzyme) | 0.00% | 0.00% |
| *C37H5.14* (Decapping enzyme) | 0.00% | 0.00% |
| *prg-1* | 74.00% | 2.49% |

**Supplementary Table 2**

List of piRNAs that show upregulation at precursor levels with the corresponding mature forms downregulated upon XRN-2 depletion compared to control (Related to Figure 2).

| **Name** | **Transcript_id** | **Gene_id** | **Sequence** |
| --- | --- | --- | --- |
| 21ur-1315 | Y51H4A.346 | WBGene00050101 | GGTTAAGAACATTTGGAAAAT |
| 21ur-13777 | C46G7.104 | WBGene00174691 | GTTTTTGAATTTCTCGGTGCT |
| 21ur-4066 | Y51H4A.325 | WBGene00049801 | GTGGTTTGATCAAAAGTGAAA |
| 21ur-4264 | Y51H4A.315 | WBGene00049588 | TTAACTTAAGTAGATAAATGA |
| 21ur-4569 | Y51H4A.118 | WBGene00046917 | CGGGTTATTGATTTACCACTA |
| 21ur-700 | Y51H4A.60 | WBGene00045998 | ATACTGTATTTGTAAGCCATG |
| piR-cel-10067 | Y105C5A.732 | WBGene00168073 | TTATTGGGCTAAACGACTGAG |
| piR-cel-1008 | R05A10.44 | WBGene00048965 | TGATGTTGTTGTGTTGTTCTT |
| piR-cel-10112 | Y105C5A.1036 | WBGene00171875 | TCTTTATAGGTTTCATTTCTG |
| piR-cel-10414 | Y40H7A.157 | WBGene00165189 | TATTGGAGGCCTGGTTGTTTG |
| piR-cel-10550 | M199.121 | WBGene00172685 | TTTATGGATTGTTCCGCATGA |
| piR-cel-10686 | T08B6.59 | WBGene00174315 | TGACGGCATAAGTTTGACATC |
| piR-cel-10700 | Y73F8A.744 | WBGene00168933 | TCTTCGTTGCAAGCTCATTTT |
| piR-cel-1077 | LLC1.19 | WBGene00047717 | TAGAACTGCTTTTATTTATTG |
| piR-cel-10859 | F13G11.136 | WBGene00174303 | TTTTATTTTCGTTTCTGTCCA |
| piR-cel-10952 | F55B11.80 | WBGene00168009 | TTGAGACTTTCCTGACTCATT |
| piR-cel-1097 | Y37A1B.109 | WBGene00048762 | TATCGCGCTCCCCGCTTCGAA |
| piR-cel-11402 | Y40H7A.174 | WBGene00166329 | TTGCCACAACAGCTCAGGGAA |
| piR-cel-11444 | Y105C5B.1208 | WBGene00172566 | TTCGAAGGTTTTTTTTTGGGA |
| piR-cel-11579 | Y57G11C.897 | WBGene00171263 | TGCCTACTACTGGAATCCATT |
| piR-cel-11676 | Y45F10B.39 | WBGene00168270 | TATTTTTTCTAGACCCATATG |
| piR-cel-11792 | Y37A1B.256 | WBGene00170544 | GACACCGATAAGAGAACAAAA |
| piR-cel-1193 | Y57G11C.448 | WBGene00050386 | TCATAACAAACTGGGCATAAA |
| piR-cel-11976 | Y67A10A.254 | WBGene00170667 | TTGTCATAATGTTGGGAGAAG |
| piR-cel-12047 | T04D1.6 | WBGene00171130 | TTCGGATCCAATATGTCACCG |
| piR-cel-12143 | Y51H4A.534 | WBGene00167342 | TTAAATCACACAGAGTAATGA |
| piR-cel-12167 | C30H6.13 | WBGene00167596 | TGTTGTTGTTGGTGTAGTGGT |
| piR-cel-12184 | T06A10.92 | WBGene00173148 | TGTTAGAATTGTCACAACCGT |
| piR-cel-12248 | Y73F8A.992 | WBGene00172663 | TGCTAGATTTTGGGTCCCATT |
| piR-cel-12503 | Y51H4A.842 | WBGene00173295 | TAATGTTGGTAAAGCTGTGGA |
| piR-cel-12707 | Y105C5B.989 | WBGene00169769 | TTTGGAACAAAGCAAATTTAA |
| piR-cel-12766 | Y57G11C.555 | WBGene00166000 | TTTCAATTGTTTAATTCAGGC |
| piR-cel-1278 | Y116A8C.150 | WBGene00049549 | TATTGACGTTTTCAGAATTGA |
| piR-cel-13022 | F55B11.131 | WBGene00174968 | TTATTTCTCCTATCAACAAGC |
| piR-cel-13060 | LLC1.80 | WBGene00170803 | TTAGGCAAGAGCGGGTGAAGA |
| piR-cel-13233 | C09B9.31 | WBGene00166303 | TGATTGGTTTCGGATTCTGCT |
| piR-cel-13323 | Y57G11C.877 | WBGene00170839 | TCTCGGATTGATTATCAGATT |
| piR-cel-13456 | Y116A8C.334 | WBGene00170155 | TATTTCGGGAATTTTCTATTT |
| piR-cel-1362 | R05A10.25 | WBGene00047075 | TGGTGCATGGCGTGGCTTTGT |
| piR-cel-13705 | Y57G11C.773 | WBGene00169290 | CGAAGTACAAATGTCAAAAAC |
| piR-cel-13755 | Y105C5A.1057 | WBGene00172143 | TTTTTTGAGTCACGGTGATTA |
| piR-cel-1377 | Y67A10A.21 | WBGene00045724 | TGCGTAAGCTAGCCGAATTGG |
| piR-cel-13777 | C46G7.97 | WBGene00172882 | TTTTTGAATTTCTCGGTGCTG |
| piR-cel-138 | Y67A10A.77 | WBGene00048206 | TCTGCAATCGATTTCTGCAAT |
| piR-cel-13971 | T27E7.70 | WBGene00172456 | TTTGAGCTCGCCGATTTCTCT |
| piR-cel-140 | Y67A10A.47 | WBGene00046763 | TGGTGTTGCCAGAAGATATAT |
| piR-cel-14117 | Y105C5B.942 | WBGene00169187 | TTTAACGTCTGAATCATGCGA |
| piR-cel-14120 | C02B10.37 | WBGene00167266 | TTTAAAGTTCTCTCGCATTTC |
| piR-cel-14189 | Y7A9C.25 | WBGene00170945 | TTGGGCAAAAGTTAGGCAGGC |
| piR-cel-14319 | C02B10.59 | WBGene00171382 | TTCTGATTTGCTCCGGTGTCA |
| piR-cel-14391 | Y59E9AL.24 | WBGene00169124 | TTCATTAGACTTGGCGTAATC |
| piR-cel-1440 | M03D4.23 | WBGene00049881 | TGATAAATTGTGCGAATTTTG |
| piR-cel-14440 | VY10G11R.70 | WBGene00174424 | TTCAAGCAAGGTTACCGAAAT |
| piR-cel-1449 | Y40H7A.78 | WBGene00047945 | TGCCACAACAGCTCAGGGAAA |
| piR-cel-14535 | Y43D4A.121 | WBGene00170348 | TTAGTATTTCTTCCTCTGATT |
| piR-cel-14655 | Y105C5A.682 | WBGene00167407 | TGTTTGTGACAGAACGCATTT |
| piR-cel-15239 | Y51H4A.570 | WBGene00168078 | TATTAAATCACACAGAGTAAT |
| piR-cel-15623 | Y41E3.261 | WBGene00166677 | CGATAAGCGTTTCAACGAAGA |
| piR-cel-15639 | C06E4.10 | WBGene00167060 | CAGGCTTTGCAGAAAGTGGTG |
| piR-cel-15645 | Y73F8A.631 | WBGene00166868 | CACATGCTTGGACTCGATAGT |
| piR-cel-1601 | C45E5.12 | WBGene00047132 | TGCTAAATGATTGATTGCTAG |
| piR-cel-1620 | Y116A8C.58 | WBGene00045762 | TACTACGATATTATTTTTTGT |
| piR-cel-1639 | Y73F8A.351 | WBGene00049167 | TGCTTGTATGTATACTTTGTG |
| piR-cel-1667 | Y57G11A.46 | WBGene00048222 | TTATGATGAGATGATTTTGGG |
| piR-cel-1714 | Y105C5A.290 | WBGene00048416 | TGGGCTTCTTGAAAACTGTAC |
| piR-cel-179 | Y43D4A.49 | WBGene00049134 | TAGTATTTCTTCCTCTGATTC |
| piR-cel-1846 | Y57G11C.482 | WBGene00050701 | TTCAATTGTTTAATTCAGGCT |
| piR-cel-1882 | Y40H7A.108 | WBGene00049253 | TTACAGTATTGGTAAGGTGGA |
| piR-cel-1894 | T23G4.8 | WBGene00045863 | TAATAAAGTACTATTGCCGTT |
| piR-cel-1943 | Y57G11B.40 | WBGene00047387 | TCAAGTCAGACAAATGTGCTT |
| piR-cel-1950 | Y105C5B.336 | WBGene00048348 | TCGAAGGTTTTTTTTTGGGAT |
| piR-cel-1954 | Y10G11A.7 | WBGene00045966 | TCAGTCTTTGAACGCAGCAAT |
| piR-cel-2024 | Y105C5A.154 | WBGene00046844 | TAAATTCGTGTCCTATTCCTA |
| piR-cel-215 | Y41E3.66 | WBGene00046889 | TAAAGTATTTGGGAACTTGGC |
| piR-cel-2268 | Y105C5A.462 | WBGene00050257 | TACTGAAGAAGACGGACAAAG |
| piR-cel-2298 | F02H6.12 | WBGene00046114 | TGTAAGCACTGTTCCCGTTTG |
| piR-cel-2440 | Y105C5A.240 | WBGene00047749 | TGAACTTTCGAACTAACCCAA |
| piR-cel-2445 | Y43D4A.44 | WBGene00048730 | TGGTATCCTCAAAAAGCTTTC |
| piR-cel-2477 | Y40H7A.128 | WBGene00049923 | TACAGTATTGGTAAGGTGGAA |
| piR-cel-2508 | Y116A8A.62 | WBGene00050543 | TCGCATTTAAAAATTTAACGA |
| piR-cel-2562 | Y43D4A.35 | WBGene00047708 | TCGCCCCCTCATGGATTTTCA |
| piR-cel-2639 | Y40H7A.52 | WBGene00046790 | TGGTATTCCAGTGAACGATTA |
| piR-cel-2773 | Y57G11C.327 | WBGene00048985 | TGGGAGGAAAATTGTTCAGAT |
| piR-cel-2862 | Y57G11C.261 | WBGene00048116 | TACAGCCCACTGCCATCTAGT |
| piR-cel-2871 | Y51H4A.85 | WBGene00046342 | TTCCTTACGAAATCTCCATAA |
| piR-cel-2908 | F52D4.7 | WBGene00048715 | TGATTTGGAATACGTTGATTT |
| piR-cel-2942 | M199.61 | WBGene00050524 | TAAATTTTCCTTCGGTACAAC |
| piR-cel-2994 | Y73F8A.79 | WBGene00046016 | TTCGCTCGTGTTGGCAGATAA |
| piR-cel-31 | Y105C5A.67 | WBGene00045860 | TGAATTTAGTGACATTGATAC |
| piR-cel-3111 | Y57G11C.305 | WBGene00048638 | TCTCCTACTCACTCTGGTTTT |
| piR-cel-3170 | Y105C5A.371 | WBGene00049231 | TGATTGACGACGTGGCTATCA |
| piR-cel-3181 | Y105C5A.276 | WBGene00048294 | TCGGCCCATCAATTTTAGCAA |
| piR-cel-3336 | M199.14 | WBGene00046062 | TATCTTCGTTTTGGCAATAAC |
| piR-cel-3520 | LLC1.21 | WBGene00047827 | TCAACTCTGATGTTTCCTTGA |
| piR-cel-3594 | Y41E3.122 | WBGene00048181 | TATTTAGTTTTCAGTATAGGC |
| piR-cel-3609 | Y105C5B.280 | WBGene00047908 | TGGAGTTTTGATTGTATGTCA |
| piR-cel-3693 | Y116A8C.144 | WBGene00049305 | TGGGTATTTGACAATTTTTGA |
| piR-cel-3911 | Y37A1A.8 | WBGene00045793 | TCGAAATTTCTTTTTGCTTCA |
| piR-cel-3996 | C06E7.8 | WBGene00045735 | TTGGATCAAAAAACAAAAACT |
| piR-cel-4018 | Y105C5A.269 | WBGene00048215 | TAAATTGCTCGATTTCACAAA |
| piR-cel-4240 | C52D10.36 | WBGene00048211 | TTGCAAACCACCACAACATTG |
| piR-cel-4377 | Y105C5B.537 | WBGene00050324 | TTTCGGAGATTCAAACACGAG |
| piR-cel-4405 | F38C2.36 | WBGene00048780 | TGTTGACTGCCTCTCTTGGAA |
| piR-cel-445 | Y105C5B.252 | WBGene00047655 | TAGTTTCTTTTTTATGTAGTC |
| piR-cel-4624 | Y73F8A.84 | WBGene00046081 | TAATATTTTTCAAGAAGGTGA |
| piR-cel-4682 | H04M03.16 | WBGene00045855 | TGATCGCACTGCTAGTTGAAA |
| piR-cel-4811 | T27E7.35 | WBGene00050078 | TGTCGAATCGCCTTCTGTAAA |
| piR-cel-4918 | Y105C5A.359 | WBGene00049085 | TACTCGGCCCGATTTATGATT |
| piR-cel-4976 | Y116A8C.110 | WBGene00047928 | TAGCGATTGCAGAGTAAGATT |
| piR-cel-5155 | Y73F8A.225 | WBGene00047763 | TACTCCTGTCCTCTCCAACAT |
| piR-cel-5215 | Y41E3.106 | WBGene00047846 | TGGTCTAATAAATTTTTGAAA |
| piR-cel-5282 | C09B9.10 | WBGene00046761 | TGCATACACGGTACTCAGAGG |
| piR-cel-5320 | H32C10.16 | WBGene00049630 | TCAGTTTTGAACAGCGGTAAC |
| piR-cel-5356 | Y67A10A.66 | WBGene00047595 | TGTTGCCAGAAGATATATAGT |
| piR-cel-5522 | Y57G11C.881 | WBGene00170993 | TTGGTGCCTACTACTGGAATC |
| piR-cel-5532 | Y116A8C.346 | WBGene00170775 | TCAGTGACAAAAAAACATCTA |
| piR-cel-5583 | Y73F8A.632 | WBGene00166929 | TATGGAAATTTGTGTTGCATT |
| piR-cel-5599 | C27H2.82 | WBGene00169024 | TTGATGTTGCACCAGGTGGAA |
| piR-cel-5608 | Y57G11C.1057 | WBGene00173763 | TAACTAGGGACAGGGCAGTTT |
| piR-cel-5700 | Y73B6BL.177 | WBGene00170393 | TGAGTATATATTGATTGTCCT |
| piR-cel-5722 | Y57G11C.591 | WBGene00166588 | TTAAAGTCACAAGAGAACAAC |
| piR-cel-5740 | Y116A8A.112 | WBGene00171901 | TTAACAGGATTAACAGGATTC |
| piR-cel-5788 | Y105C5A.878 | WBGene00169794 | TCTTGCCGGTATTGATGTTGA |
| piR-cel-581 | Y67A10A.68 | WBGene00047644 | TGTCATAATGTTGGGAGAAGT |
| piR-cel-5818 | Y105C5A.529 | WBGene00165278 | TTCCTAGGATCGTTGCACAAA |
| piR-cel-5833 | Y105C5B.1063 | WBGene00170657 | TTTGGCCTAGAAATTTAAGAT |
| piR-cel-5861 | Y51H4A.497 | WBGene00166709 | TAGAAAATATTGATTGAATCG |
| piR-cel-5913 | Y105C5A.1215 | WBGene00174235 | TGCATGGGCGGATTTCAGTTT |
| piR-cel-6031 | Y51H4A.439 | WBGene00165751 | TTTGATGTGAGTATAGAAAGT |
| piR-cel-6037 | Y57G11C.937 | WBGene00171907 | TGCTAATTGTGAACTCAATTC |
| piR-cel-6098 | Y105C5B.675 | WBGene00165950 | TTCGATTTTGAGAAAACTGGC |
| piR-cel-6262 | Y116A8B.48 | WBGene00168052 | TGGGCGTACAGTTTGAACTTC |
| piR-cel-6495 | Y41E3.225 | WBGene00165320 | TCCTCTGTCTGTTTTTCCTCG |
| piR-cel-6744 | Y105C5B.773 | WBGene00167284 | TTGGAACAAAGCAAATTTAAG |
| piR-cel-6828 | Y51H4A.758 | WBGene00171747 | TCCGCGGTCGTACGGCAGATT |
| piR-cel-7045 | Y37A1B.160 | WBGene00165580 | TATTTAATTTTGCGAGTAGAA |
| piR-cel-7061 | Y105C5B.1283 | WBGene00173523 | TTATGATTGCTTAGTGGAAGA |
| piR-cel-7497 | Y67A10A.250 | WBGene00170430 | TTATTTGAACATATACCTCAT |
| piR-cel-7611 | Y116A8A.139 | WBGene00174291 | GCATTTAAAAATTTAACGACT |
| piR-cel-7670 | Y10G11A.89 | WBGene00174898 | TACTGACTGAATTTTTTTTCT |
| piR-cel-769 | Y116A8C.73 | WBGene00046400 | TTAAATCACTGTGGTATCTAT |
| piR-cel-7705 | Y105C5A.774 | WBGene00168647 | TCTTTCAAATTATCGTTCTTT |
| piR-cel-7733 | Y105C5B.1270 | WBGene00173325 | TTTTTTATGGGTGGTCATTGG |
| piR-cel-7786 | Y105C5B.760 | WBGene00167011 | TTGTGACTCAATTTCAAGATT |
| piR-cel-7870 | Y73F8A.904 | WBGene00171313 | TTTTGCCTGTAAAAAATTTTC |
| piR-cel-792 | Y105C5B.556 | WBGene00050433 | TTATGATCTGACAACTCGTTT |
| piR-cel-type2-1134 |  | Type_II_1134 | TAATGACCGCCGGGAGCAGTT |
| piR-cel-type2-852 |  | Type_II_852 | TGAGCCGTCGTATCCCTGGAC |
| piR-cel-type2-89 |  | Type_II_89 | TTTCAGCTCTTTCAGTTCTCT |
| piR-cel-type2-898 |  | Type_II_898 | TAGGCTTCGCAGTAGATCAAG |

**Supplementary Table 3**

Endoribonucleases employed in piRNA derepression screen (Related to Figure 4).

| **Name** | **Mean** | **SEM** |
| --- | --- | --- |
| *endu-1* | 17.00% | 1.23% |
| *endu-2* | 0.00% | 0.00% |
| *hoe-1* | 2.99% | 0.34% |
| *rege-1* | 1.67% | 0.80% |
| *pelo-1* | 1.82% | 0.33% |
| *K07D4.2* | 0.00% | 0.00% |
| *Y6B3B.9* | 0.00% | 0.00% |
| *F14F9.5/B6* | 0.00% | 0.00% |
| *F44E5.15* | 0.00% | 0.00% |
| *tsen-2* | 0.00% | 0.00% |
| *tsen-34* | 0.00% | 0.00% |
| *tsen-54* | 0.00% | 0.00% |

Strains used in this study.

**Supplementary Table 4**

| **Name** | **Genotype** |
| --- | --- |
| N2 | Wild-type |
| SX1316 | *mjIs144 II; unc-119(ed3) III* |
| SX922 | *prg-1(n4357) I* |
| RB1621 | *K10C8.1(ok1994) V* |
| *xrn-2(PHX25)* | *xrn-2(D86-A, D181-A)* |

**Supplementary Table 5**

| **Cloning primers** | | |
| --- | --- | --- |
| **Gene name** | **Primer sequence (5'-3')** | **Reference** |
| *prg-1* FP | GAACCATGGCATCTGGAAGTGGTCG | This study |
| *prg-1* RP | GAAACTCGAGTTACAAGAAGAACAGCTTGTCACG | This study |
| *parn-1* FP | GTACTCGAGATGGTTATCGTCAC | This study |
| *parn-1* RP | GCCTCTAGATTATATTTCCATCGC | This study |
| *endu-1* FP | CGATCTAGAATGTGCTCTGTGCTCATTTTTATCTCAATC | This study |
| *endu-1* RP | TACCTCGAGTTAATTGAATCGATTTTGAGCTCTACACTG | This study |
| *dcap1* FP | GCATCTAGAGAACCAGRATGAGCGACG | This study |
| *dcap1* RP | GTTCCATGGTTAATCGATATTTAGACGCCGATTG | This study |
| *dcap2* FP | GTACCCGGGATGGAAATTTCTACTGAAAATTGGTGC | This study |
| *dcap2* RP | GTCTCTAGATTATGGTAATTGTGGTCCAGTGCC | This study |
| *C37H5.14* FP | GCATCTAGAATGTCAGTGAGTTTCAAGCTGGC | This study |
| *C37H5.14* RP | ATCAAGCTTATACGCGAAGAAAGTTCTGTGGC | This study |
| *eol-1* FP | ATCCTCGAGATGGCAGTTCAAGTCAAATTCGAAC | This study |
| *eol-1* RP | GCATCTAGATCACCGTTCGAAATACTCCAGG | This study |
| *FP = Forward primer, RP = Reverse primer. | | |
| **qPCR primers** | | |
| **Target gene name** | **Primer sequence (5'-3')** | **Reference** |
| *actin-1* FP | CACGGTATCGTCACCAACTG | 4 |
| *actin-1* RP | GTACGTCCGGAAGCGTAGAG | 4 |
| *Tc1* FP | AACCGTTAAGCATGGAGGTG | 4 |
| *Tc1* RP | CACATGACGACGTTGAAACC | 4 |
| *Tc3* FP | GAGCGTTCACGGAGAAGAAG | 4 |
| *Tc3* RP | AATAGTCGCGGGTTGAGTTG | 4 |
| *turmoil1* FP | AGAGACGAATGCCGTGCTAC | 9 |
| *turmoil1* RP | CGAGTCGGACACAATCAATG | 9 |
| *bath-45* FP | ACGATAGGCCTTTTCCAATG | 9 |
| *bath-45* RP | GAGCTGCCTTTTCCAAATTC | 9 |
| *xol-1* FP | GGAGCGAAAACAGTCCAGTC | 13 |
| *xol-1* RP | ATCATGTGCATGCTTCGGTA | 13 |
| *gfp spliced transcript* FP | CAAGACACGTGCTGAAGTCAA | 9 |
| *gfp spliced transcript* RP | TGTGTCCAAGAATGTTTCCATC | 9 |
| *gfp primary transcript* FP | TCTGTCAGTGGAGAGGGTGA | 9 |
| *gfp primary transcript* RP | TTTAAACTTACCCATGGAACAGG | 9 |
| *FP = Forward primer, RP = Reverse primer. | |  |
| **Northern probes** | | |
| **Target RNA** | **Sequence (5'-3')** | **Reference** |
| 21UR-1 | TGCACGGTTAACGTACGTACCAT | 17 |
| 21UR-1949 | TCCATTCTTTGTCACCCTCGTAT | 17 |
| 21UR-3372 | CTTCAATCAACTTTAGATGCA | This study |
| tRNAGly | GCTTGGAAGGCATCCATGCTGACCATT | 24 |
| 5.8S rRNA | CAACCCTGAACCAGACGTACCAACTGGAGGCCCAGTTGGT | 24 |
| 26S rRNA | CGGTACTTGTTCGCTATCGCAATCGAGTCGATATTTAGC | 24 |
| 18S rRNA | GGTTCACCTACAGCTACCTTGTTACGACTTTTACCCG | 24 |
